# Profiling extracellular vesicles in circulation enables the early detection of ovarian cancer

**DOI:** 10.1101/2023.01.19.524549

**Authors:** Ala Jo, Allen Green, Jamie E. Medina, Sonia Iyer, Anders W. Ohman, Eric T. McCarthy, Ferenc Reinhardt, Thomas Gerton, Daniel Demehin, Ranjan Mishra, David L. Kolin, Hui Zheng, Christopher P. Crum, Robert A. Weinberg, Bo R. Rueda, Cesar M. Castro, Daniela M. Dinulescu, Hahko Lee

**Author notes:** Co-Corresponding Authors (C.M.C., D.M.D., H.L.). Equal contribution (J.E.M., S.I., A.W.O., E.T.M.). Equal contribution (Cesar M. Castro, Daniela M. Dinulescu, Hahko Lee).

## Abstract

Ovarian cancer is a heterogeneous group of tumors in both cell type and natural history. While outcomes are generally favorable when detected early, the most common subtype, high-grade serous carcinoma (HGSOC), typically presents at an advanced stage and portends less favorable prognoses. Its aggressive nature has thwarted early detection efforts through conventional detection methods such as serum CA125 and ultrasound screening and thus inspired the investigation of novel biomarkers. Here, we report the systematic development of an extracellular-vesicle (EV)-based test to detect early-stage HGSOC. Our study is based on emerging insights into HGSOC biology, notably that it arises from precursor lesions within the fallopian tube before traveling to ovarian and/or peritoneal surfaces. To identify HGSOC marker candidates, we established murine fallopian tube (mFT) cells with oncogenic mutations in *Brca1/2, Tp53*, and *Pten* genes, and performed proteomic analyses on mFT EVs. The identified markers were then evaluated with an orthotopic HGSOC animal model. In serially-drawn blood samples of tumor-bearing mice, mFT-EV markers increased with tumor initiation, supporting their potential use in early cancer detection. A pilot human clinical study (*n* = 51) further narrowed EV markers to five candidates, EpCAM, CD24, VCAN, HE4, and TNC. Combined expression of these markers achieved high OvCa diagnostic accuracy (cancer vs. non-cancer) with a sensitivity of 0.89 and specificity of 0.93. The same five markers were also effective in a three-group classification: non-cancer, early-stage (I & II) HGSOC, and late-stage (III & IV) HGSOC. In particular, they differentiated early-stage HGSOC from the rest with a specificity of 0.91. Minimally invasive and repeatable, this EV-based testing could be a versatile and serial tool for informing patient care and monitoring women at high risk for ovarian cancer.

## INTRODUCTION

Ovarian cancer (OvCa) is the most lethal gynecological disease in the United States and the fifth leading cause of all cancer-related deaths among women^1^. The early-stage disease remains mostly asymptomatic, and most patients are diagnosed at late and often incurable stages. In cases where cancer is detected early, five-year survival rates exceed 90%^2^. Unfortunately, conventional screening with blood testing (e.g., CA125) and imaging have failed to demonstrate survival advantages in a large, multi-year screening trial^3^. This failure has largely been attributed to the fact that the most common OvCa subtype, high-grade serous cancer (HGSOC), often presents after spreading beyond the primary site of origin. This fact underscores the need for improved detection methods and more informed perspectives on serous carcinogenesis, tumor evolution, and metastasis^3–5^.

Amassing biological and clinical data support that the bulk of HGSOC arises from precursor lesions within the distal fallopian tube (FT)^6–12^. Patients with advanced HGSOC frequently harbor serous tubal intraepithelial carcinoma (STIC) lesions with identical *TP53* mutations found in tumors. Statistical studies indicate that nearly 60% of epithelial HGSOCs are tubal in origin^6–8,13,14^. This mechanistic insight raises the prospect of early HGSOC diagnostics by interrogating molecular markers derived from FT precursor lesions. An appealing target is extracellular vesicles (EVs) secreted by cells^15,16^. EVs reflect the molecular cargo of tumor cells and circulate in easily accessible bodily fluids^17–21^. Analyzing EVs can thus represent a real-time, minimally invasive modality to detect and monitor tumor molecular status, including precursor lesions such as STICs^22–24^. Already, tumor-associated EVs have been demonstrated to be effective surrogate OvCa biomarkers for tumor detection and treatment monitoring^25,26^. Most studies, however, mainly analyzed late-stage diseases, skewing EV markers towards advanced clinical presentations^23,26–28^. Conceivably, early OvCa lesions may have molecular profiles distinct from those of late-stage diseases or cell cultures – identifying and validating such EV signatures is a crucial to improving early diagnosis^29^.

Here, we report on our systematic approach to developing an EV-based blood test for early-stage (or low-volume) OvCa detection. We specifically tested the hypothesis that EVs from precursor lesions can be identified in blood to serve as circulating biomarkers for pre-invasive / early-stage HGSOC (**Fig. 1**). The study began by establishing murine fallopian tube (mFT)-derived tumor cell lines; these cells carried the genetic alterations (i.e., *Brca, Tp53, Pten*) commonly observed in STICs and HGSOC^30^ and recapitulated tumor progression when implanted orthotopically. We next identified mFT-specific EV markers through proteomic profiling and monitored them in serially-drawn mouse blood samples. A high-throughput assay (384-well format, 50 nL plasma per marker) with signal amplification was employed to reliably detect low-level EVs. The assay revealed that circulating mFT-EV markers increased with tumor initiation, supporting their potential use in early cancer detection. In the ensuing pilot clinical study (*n* = 51), we further refined the FT-EV signature (CD24, EpCAM, HE4, TNC, VCAN) to achieve high OvCa diagnostic accuracy (cancer vs. non-cancer) with a sensitivity of 0.89 and specificity of 0.93. The same five markers also effectively classified three clinically distinct groups: non-cancer (*n* = 14), early-stage HGSOC (stage I, II; *n* = 17), and late-stage HGSOC (stage III, IV; *n* = 20). In particular, they differentiated early-stage HGSOC from the rest with a specificity of 0.91 (= 31/34). Overall, our EV-based liquid biopsy could help reduce the high prevalence of late-stage HGSOC diagnoses or effectively monitor high-risk women.

**Figure 1.**
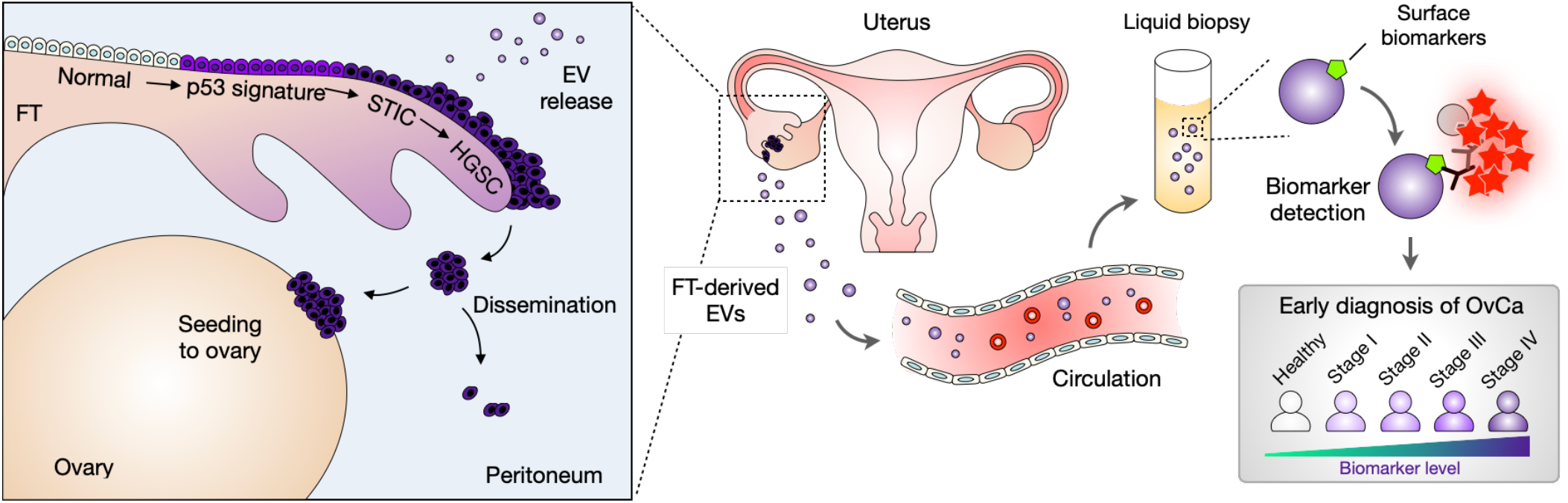
Study design. High-grade serous ovarian cancer (HGSOC) is considered to arise from precursor lesions within the fallopian tube (FT). Circulating EVs from FT precursor lesions thus can serve as early HGSOC biomarkers. In this study, we identified EV markers specific to FT carcinoma and detected them in blood samples of OvCa patients. Analyzing surface markers on FT-derived EVs allowed for differentiating early- (stage I & II) and late-stage (stage III & IV) OvCa patients.

## RESULTS

### Generating murine fallopian tube-derived tumor cell lines

We established murine fallopian tube (mFT) tumor cell lines from genetically engineered mouse models (GEMMs) of HGSOC^31^ (**Fig. 2A**). The GEMMs contained a *Pax8-Cre* transgene and different combinations of *Brca* (*Brca1* or *Brca2*), *Tp53*, and *Pten* floxed genes. Under these constructs, Cre-recombinase expression can be driven by the *Pax8* promoter; PAX8 is a transcription factor specific to Müllerian-derived epithelia (e.g., FT) but not the ovaries^32^. We isolated non-induced mFT cells from GEMM cohorts and treated the cells with doxycycline, inactivating target genes *via* Cre-mediated recombination. These processes produced oncogenic mFT cell lines with different genotypes: mFT3707 (*Brca1*^*+/–*^, *Tp53*^*mut*^, *Pten*^*–/–*^), mFT3635 (*Brca1*^*–/–*^, *Tp53*^*mut*^, *Pten*^*–/–*^), mFT3665 (*Brca2*^*+/–*^, *Tp53*^*mut*^, *Pten*^*–/–*^), and mFT3666 (*Brca2*^*–/–*^, *Tp53*^*mut*^, *Pten*^*-/-*^). We further transfected the cell lines with mCherry/luciferase plasmid and sorted them according to mCherry expression (see **Methods** for details).

**Figure 2.**
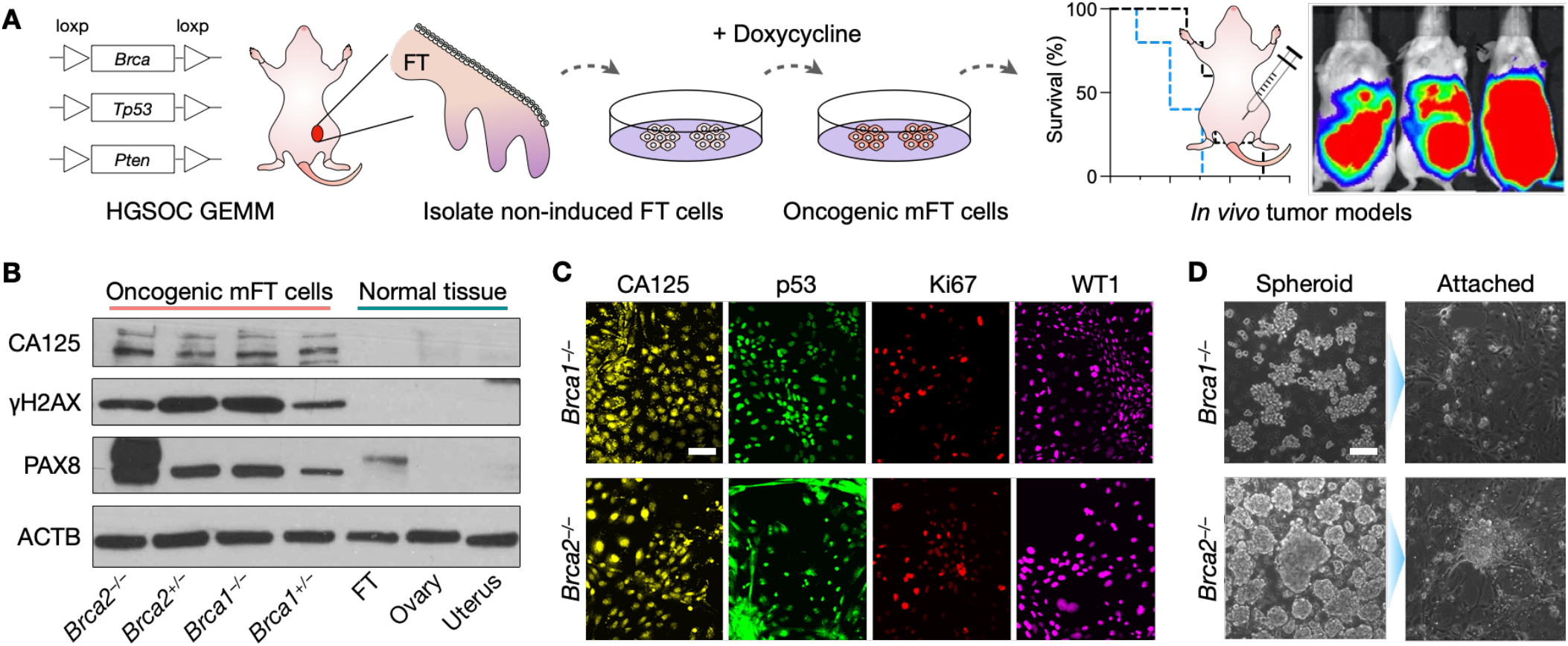
Generation and characterization of mFT cell lines. **(A)** FT cells were isolated from genetically engineered mice harboring mutations in *Brca1* or *Brca2*, as well as *Tp53* and *Pten*. Isolated cells were rendered oncogenic through the doxycycline treatment. Tumor animal models were generated by implanting the transformed cells into mice. GEMM, genetically engineered mouse model. **(B)** Oncogenic mFT cells expressed FT-specific protein (PAX8) and HGSOC markers (CA125, γH2AX). Normal tissue samples (uterus, ovary, FT) were devoid of HGSOC markers, while PAX8 was positive only with FT tissue. **(C)** Immunofluorescence microscopy confirmed that the oncogenic mFT cells (mFT3635 and mFT3666) expressed key HGSOC markers (CA125, p53, Ki67, WT1) at the cellular level. Scale bar, 50 µm. **(D)** Under *in-vitro* ultra-low adherence culture conditions, oncogenic mFT cells (mFT3635 and mFT3666) formed tumor spheroids (left). When transferred to adhesion plates, tumor spheroids adhered to the surface and spread, demonstrating their capacity to engraft. Scale bar, 100 µm.

The transformed mFT cell lines had their intended *Brca, Pten*, and *Tp53* genotypes verified by targeted polymerase chain reactions (**Fig. S1**). At the protein level, the mFT cell lines maintained the expression of the FT epithelial marker (PAX8), but they acquired *de novo* expression of HGSOC markers (CA125, γ-H2AX) that were absent in normal uterine, ovarian, and FT tissues^33,34^(**Fig. 2B**). Immunofluorescence imaging further confirmed that the mFT cell lines expressed key HGSOC markers (CA125, TP53, Ki67, WT1; **Fig. 2C** and **Fig. S2A**)^34–36^. When cultured *in vitro* on ultra-low adhesion plates, the transformed mFT cells formed spheroids. When these spheroids were re-introduced to adherent conditions, they bound to a surface and formed a monolayer, demonstrating their potential to engraft and grow into tumors (**Fig. 2D** and **Fig. S2B**).

### Characterization of EVs from parental mFT cell lines

We next analyzed mFT cell-derived EVs (mFT-EVs). Transformed mFT cell lines with two distinct genotypes, mFT3635 (*Brca1*^*–/–*^) and mFT3666 (*Brca2*^*–/–*^), were cultured in identical conditions, and vesicles in the culture media were collected *via* size exclusion chromatography (see **Methods**). We observed no significant difference in physical profiles between these two sample types. Vesicles displayed a similar morphology under electron microscopy (**Fig. 3A** and **Fig. S3**), with a size range of 50 – 150 nm (**Fig. 3B**). Nanoparticle tracking analysis also showed that samples from both cell types had a similar size distribution and concentration (**Fig. 3C**).

**Figure 3.**
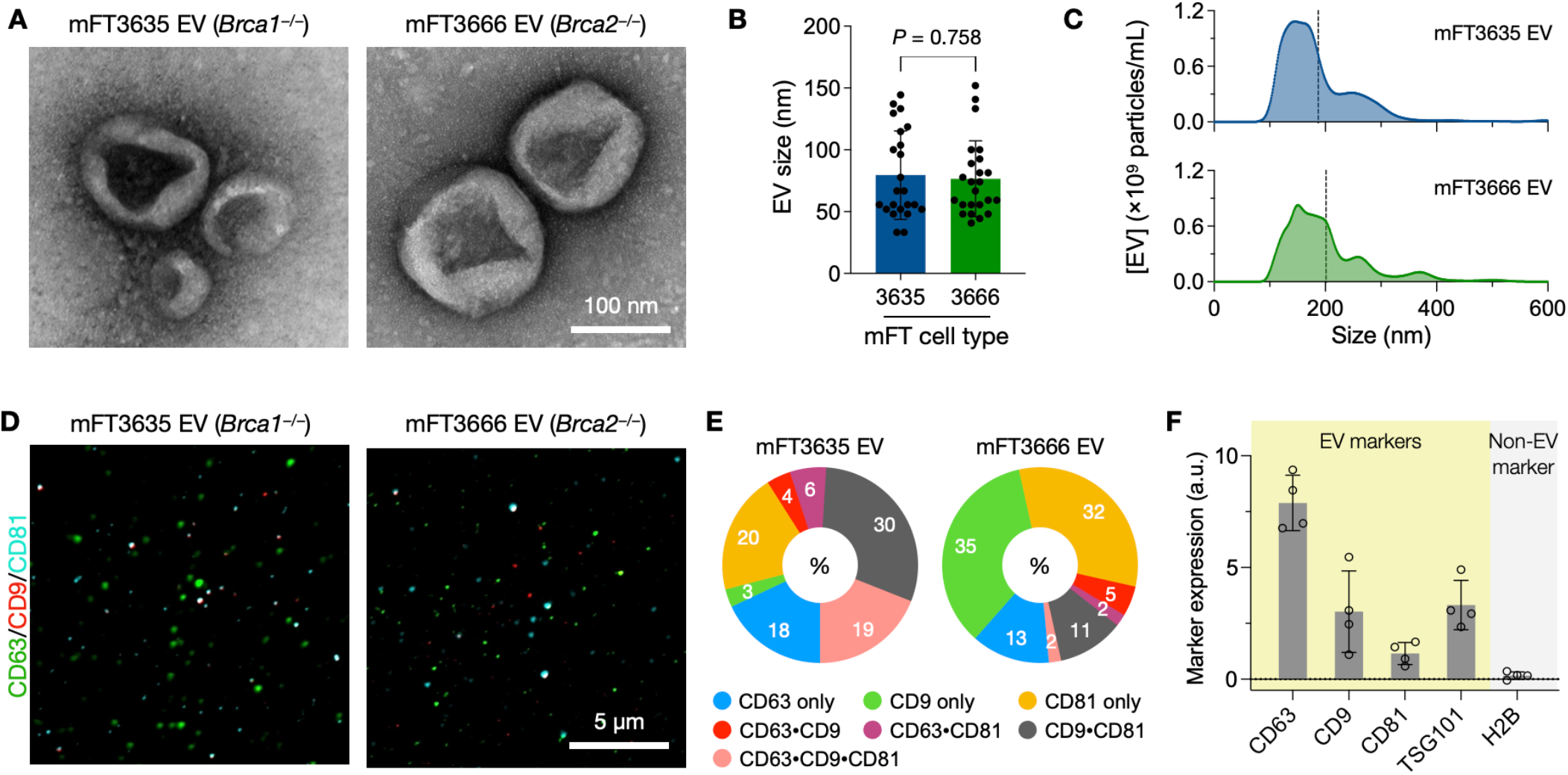
Characterization of EVs from mFT cell lines. **(A)** Transmission electron microscopy (TEM) images of EVs from oncogenic mFT cells. **(B)** Size distribution of EVs in TEM images. No significant difference (*P* = 0.758, unpaired two-sided *t-*test) was observed. Each dot represents a single EV. Error bars, s.d. **(C)** Nanoparticle tracking analysis of mFT EVs. Repeated measurements (*n* = 3) were combined. EVs from both mFT cell lines had a similar distribution of the hydrodynamic diameter with a mean value of 187 nm (mFT3635) and 201 nm (mFT3666). **(D)** Single EV imaging detected the presence of tetraspanins, CD63, CD9, and CD81, in the mFT EVs. Note that EVs appeared larger than their actual size due to optical diffraction. **(E)** Profile of tetraspanin expression from single EV imaging. Most EVs expressed only one type of tetraspanin. **(F)** Bulk EV analysis (ELISA) confirmed that mFT EVs were enriched with canonical EV markers (*i*.*e*., CD63, CD9, CD81, TSG101) and devoid of a non-EV marker (histone H2B), a.u., arbitrary units. Data are displayed as mean ± s.d. (*n* = 4).

Single particle imaging confirmed vesicular identity. We captured vesicles on a glass surface and immunolabeled them for tetraspanins (CD63, CD9, CD81) enriched in EVs (**Fig. 3D**). A large portion (41% for mFT3635 EV and 80% for mFT3666 EV) of vesicles expressed only one type of EV-related tetraspanin (**Fig. 3E**). The results indicated that relying on single tetraspanins may underestimate EV numbers and lower analytical sensitivity. A separate bulk assay (ELISA) further confirmed EV’s presence (**Fig. 3F**): the samples were positive for canonical EV markers (i.e., CD63, CD9, CD81, TSG101) and devoid of a non-EV marker (i.e., histone H2B).

### mFT-EV marker selection

To determine protein candidates from the mFT tumor, we carried out comprehensive EV proteomic analyses and processed the results through a bioinformatic pipeline (**Fig. 4A**). EVs were collected from mFT3635 (*Brca1*^*–/–*^) and mFT3666 (*Brca2*^*–/–*^) cells and subjected to liquid chromatography-mass spectrometry. The process identified 677 proteins from six replicates (see **Supplementary Data**): 631 proteins from mFT3635-EVs and 215 proteins from mFT3666-EVs (**Fig. S4**). We selected 169 proteins found in both EV types (**Fig. 4B**), effectively identifying markers relevant to both *Brca1/2* mutations. We then further filtered the list through public databases: i) cell proteome data (UniProt) for membrane-associated proteins and ii) an established EV proteome database (EVpedia) for markers present in EVs. We finally augmented the list with additional protein markers from the literature.

**Figure 4.**
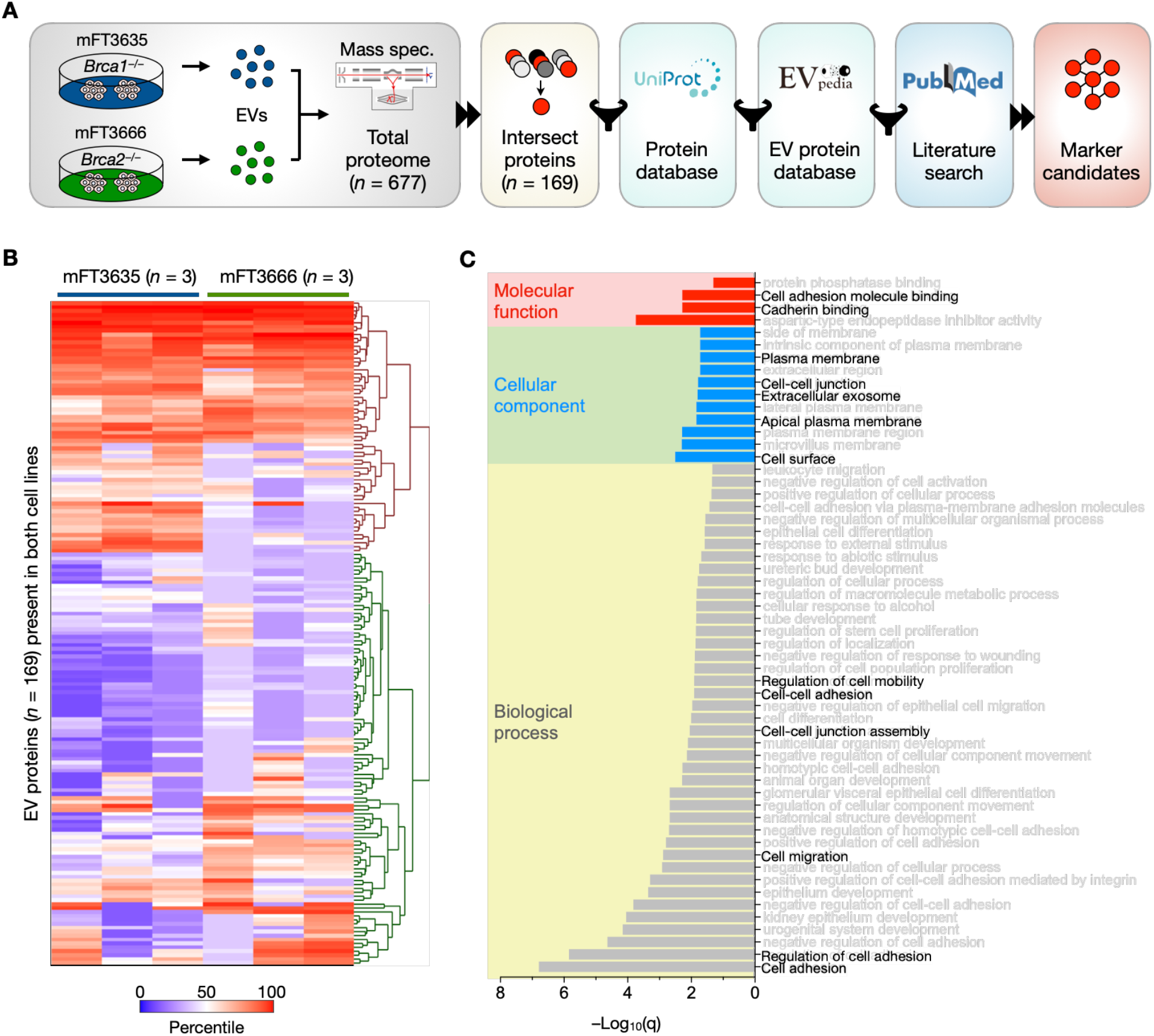
mFT-EV marker selection. **(A)** Selection algorithm. EVs from mFT3635 and mFT3666 cells were isolated and processed for proteomic analysis. Acquired data were filtered through public databases for their location in the cell membrane (Uniprot) and presence in EVs (EVpedia), and the outcomes were further curated through a literature search (PubMed). **(B)** Heatmap of proteins (*n* = 169) found in EVs from mFT3635 and mFT3666 cell lines. The data-driven approach selected nine candidate markers (PODXL, JUP, TNC, VCAN, CD24, EpCAM, HE4, FOLR1, CA125) from highly expressed proteins. **(C)** Gene ontology (GO) analysis showed that the selected markers were strongly associated with cellular adhesion. GO analysis was performed with STRING v11.5.

The selection algorithm produced the final nine candidate markers: podocalyxin (PODXL), junction plakoglobin (JUP), tenascin-C (TNC), versican (VCAN), CD24, epithelial cell adhesion molecule (EpCAM), human epididymis protein 4 (HE4), folate receptor 1 (FOLR1), cancer antigen 125 (CA125). Gene ontology analysis confirmed the enrichment of these markers in the plasma membrane and EVs (**Fig. 4C**). The dominant function was cellular adhesion (PODXL, JUP, TNC, VCAN, CD24, EpCAM)^37– 42^, followed by immune responses within the cell (HE4, CA125)^43,44^ and DNA repair (FOLR1)^45^. We further validated the expression of candidate markers in mFT parental cells (**Fig. S5A**). All markers stained positive in oncogenic mFT cell lines. Interestingly, these markers were also positive in cells collected from OvCa mice ascites (**Fig. S5B**), suggesting that they may be expressed throughout disease progression or peritoneal washings.

### Sensitive, high-throughput EV assay

We next established an EV assay with superior sensitivity and throughput, which was critical to detecting multiple proteins in low-abundant EVs from early cancer lesions; **Figure 5A** summarizes our assay scheme. Adopting the reverse-phase protein array, we directly immobilized EVs on microwell plate surfaces through physical adsorption (pH = 9.4)^46^; this step allowed for unbiased EV capture regardless of tetraspanin expression. We then applied tyramide signal amplification (TSA) to boost the detection sensitivity. Target EV proteins were labeled with primary antibodies, which were further labeled with horseradish peroxidase (HRP) through secondary antibodies. When tyramide-biotin and H_2_O_2_ were added, HRP-labeled EVs catalyzed the production of reactive tyramide radicals, triggering the dense deposition of biotin molecules on nearby tyrosine residues^47^. Finally, fluorescent streptavidin was coupled to the biotin deposit for signal generation.

**Figure 5.**
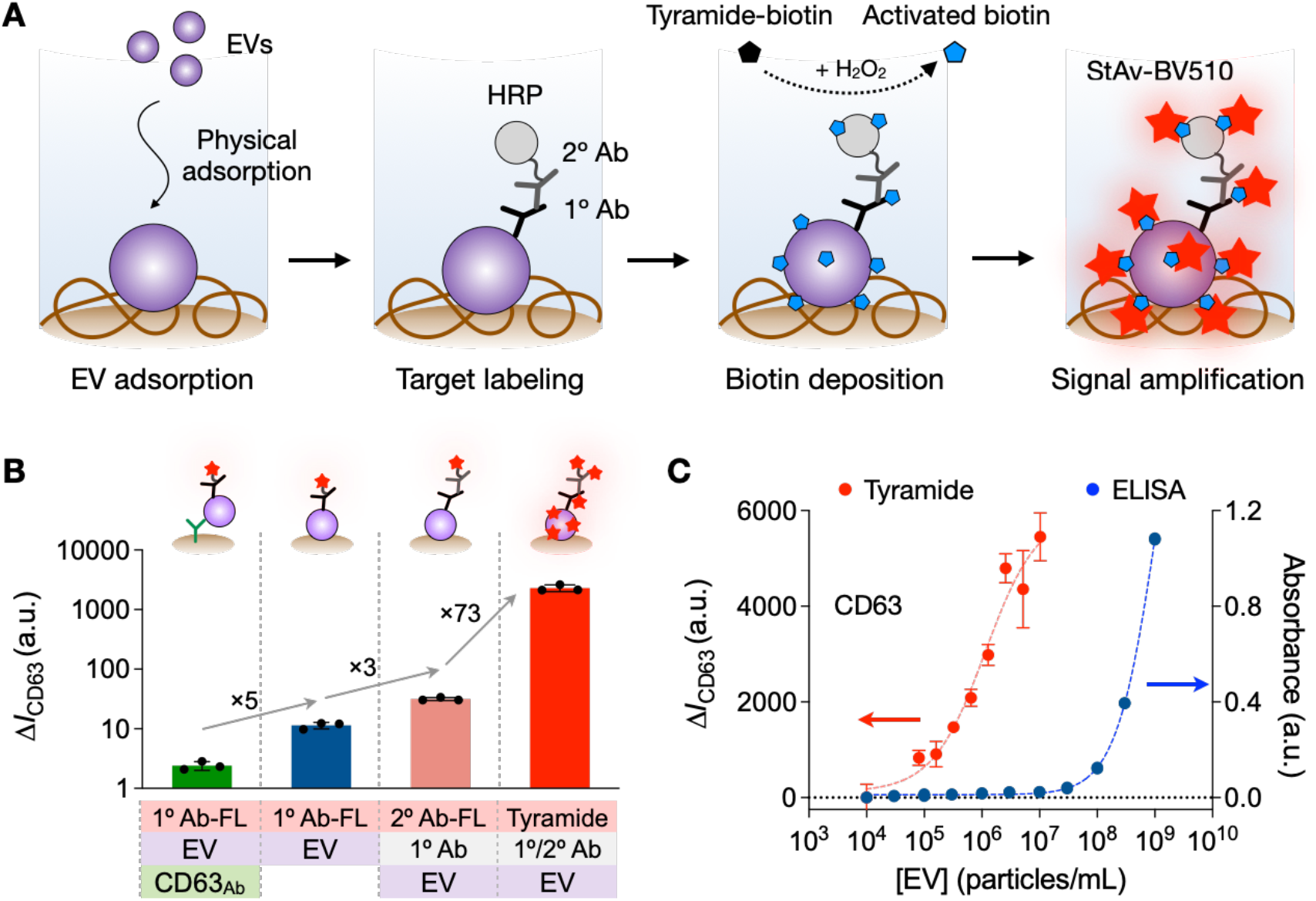
Sensitive, high-throughput EV assay. **(A)** Assay scheme. EVs are captured on a polystyrene plate *via* physical adsorption. Target EV protein is labeled with a primary antibody (1° Ab) which is further labeled with a secondary antibody (2° Ab) conjugated with horseradish peroxidase (HRP). With tyramide-biotin and hydrogen peroxide (H_2_O_2_) added, the HRP catalyzes the dense deposition of biotin. Finally, the analytical signal is detected by adding fluorescent streptavidin (StAv-BV510). **(B)** Different EV-assay formats were compared for the analytical signal. Physisorptive EV immobilization produced a higher signal (5-fold) than Ab-based EV capture (left). Among the physisorptive EV assays, applying the tyramide amplification generated the highest signal. The overall signal increase was about 10^3^-fold. Data are displayed as mean ± s.d. (*n* = 3). FL, fluorescent. **(C)** The tyramide-amplification assay (TSA) displayed superior sensitivity compared to conventional ELISA. Based on CD63 titration curves, the estimated detection limits were 2.4 × 10^4^ EV/mL for TSA (6 × 10^2^ EVs in 25 μL) and 3 × 10^8^ EV/mL for ELISA (1.5 × 10^6^ EVs in 50 μL). a.u., arbitrary units.

The developed TSA assay reported the highest analytical signal among the different assay configurations tested (**Fig. 5B**). As a model system, we probed mFT3635 EVs for CD63 expression, measuring the net fluorescence intensity, Δ*I*_CD63_ = *I*_CD63_ – *I*_IgG_, wherein *I*_CD63_ was the signal from samples labeled with CD63 antibody and *I*_IgG_ was from samples labeled with an isotype IgG antibody. Comparing EV-immobilization results, we observed a 5-fold signal increase when switching from the antibody-based EV capture to EV physisorption. The signal further improved (3-fold) when a fluorescent secondary antibody was used for labeling. Peak intensity was reached when the aforementioned TSA method was employed: a 73-fold increase over the next best detection method.

The large signal gain led to superb analytical sensitivity. Varying the input EV loading, we measured Δ*I*_CD63_ (**Fig. 5C**). The TSA assay achieved a limit of detection (6 × 10^2^ EVs) that was much lower than conventional sandwich-type ELISA (1.5 × 10^6^ EVs). Such high sensitivity allowed the TSA assay to detect markers within a small volume (50 nL) of plasma, which allowed us to use a 384-well plate to increase overall throughput. The TSA assay was also robust, displaying high correlations in the inter-assay comparison (**Fig. S6**).

### Serial profiling of plasma EVs from mFT-tumor mice

We next performed a longitudinal mouse study evaluating EVs for early OvCa detection (**Fig. 6A**). To replicate mFT tumor initiation and progression in the ovary, we implanted oncogenic mFT cells into the ovarian fat pad/bursa of NSG mice. Both mFT cell types, mFT3635 (*Brca1*^*–/–*^) and mFT3666 (*Brca2*^*–/–*^), developed into a tumor with no significant difference (*P* = 0.817; log-rank test) in survival rates (**Fig. 6B**). When these cell lines were implanted intraperitoneally (IP) into NOD SCID mice, the tumor spread throughout the peritoneal cavity and caused extensive ascites (**Fig. S7**), recapitulating *in situ* and late-stage diseases. Immunohistochemical staining revealed that the late-stage tumor (IP-engrafted) was positive for PAX8 (FT epithelial marker)^32^, TP53 (HGSOC marker)^35^, WT1 (HGSOC marker)^36^, and STMN1 (tumor progression)^48^, confirming the tumor’s mFT origin and metastatic potential (**Fig. 6C**).

**Figure 6.**
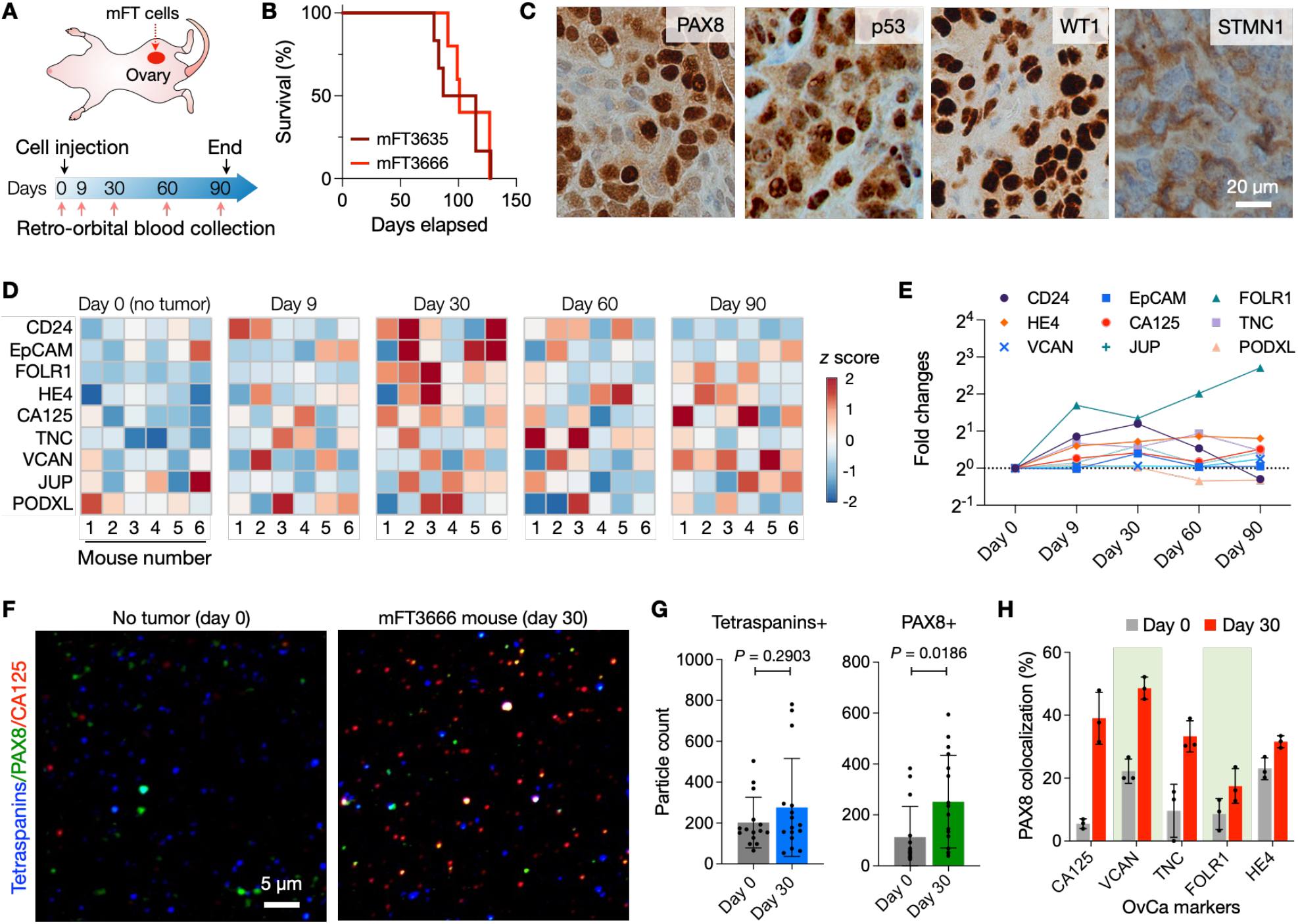
Serial EV profiling with an OvCa animal model. **(A)** Study design. Oncogenic mFT cells were implanted into ovary fat mass in mice. Blood samples were collected from the animals before engraftment and up to three months thereafter. For each sample, EVs were collected and profiled for nine candidate markers. **(B)** Survival analysis of mFT3635 (*Brca1*^−/–^) and mFT3666 (*Brca2*^−/–^) implanted animals. No significant difference (*P* = 0.989; log-rank test) was observed between survival curves. The median survival was 107 days (mFT3635, *n* = 6) and 114 days (mFT3666, *n* = 6). **(C)** Immunohistochemical staining confirmed the expression of the FT epithelial (PAX8) and tumor markers (p53, WT1, STMN1) in an IP-engrafted late-stage (day 90) tumor. **(D)** Longitudinal EV profiling in tumor-bearing animals (*n* = 6). Nine OvCa markers were measured in plasma EVs. The z-score of each marker was used to generate the heatmap. **(E)** The overall expression of OvCa markers increased after tumor initiation (day 9) and peaked 30 days after the mFT cell implant. **(F)** Single EVs were imaged in plasma samples collected before the mFT engraftment (day 0) and during disease progression (day 30). EVs were stained for tetraspanins (CD63, CD9), PAX8 (FT epithelial marker), and CA125 marker (see **Fig. S8** for other OvCa markers). **(G)** Tetraspanin-positive EVs were present in both samples (day 0 and day 30) with no significant difference in numbers (*P* = 0.2903; non-paired, two-sided *t*-test). PAX8-positive EVs, however, significantly increased in the tumor sample (*P* = 0.0186; non-paired, two-sided *t*-test). Data are displayed as mean ± s.d. (*n* = 15 field of views). **(H)** EV imaging revealed that more EVs were both PAX8 and OvCa-marker positive in tumor samples. Data are displayed as mean ± s.d. (*n* = 3). Non-paired, two-sided *t*-test: *P* = 0.002 (CA125); *P* = 0.0009 (VCAN); *P* = 0.014 (TNC); *P* = 0.105 (FOLR1); *P* = 0.018 (HE4).

For longitudinal EV profiling, we collected serial blood samples from the host animals (*n* = 6) starting before engraftment and continuing up to 3 months thereafter. Our high-throughput TSA assay was then applied to detect mFT markers in plasma EVs (**Fig. 6D**; see **Methods** for details). The overall expression of mFT markers elevated in all animals after tumor initiation (Day 9) and peaked 30 days after the mFT cell implant (**Fig. 6E**), which supported EVs’ potential for early cancer detection.

We further performed single EV imaging of plasma samples collected before mFT-cell implantation (Day 0) and during tumor growth (Day 30). Samples were stained for tetraspanins (CD63, CD9; EV identification), an FT epithelial marker (PAX8), and OvCa markers (**Fig. 6F**). No significant changes were observed in tetraspanin-positive EV numbers between no-tumor (Day 0) and tumor (Day 30) samples, whereas PAX8-positive EV numbers significantly increased in the tumor samples (**Fig. 6G**). Notably, more EVs were PAX8 and OvCa-marker positive in tumor samples (**Fig. 6H** and **Fig. S8**), validating the presence of FT-derived tumor EVs in circulation.

### Profiling plasma EVs from clinical samples

We conducted a pilot study evaluating the EV-marker candidates’ performance in human clinical samples (**Table 1**). Plasma samples were obtained from HGSOC patients (*n* = 37) at different stages (*n* = 7, Stage I; *n* = 10, Stage II; *n* = 10, Stage III; *n* = 10, Stage IV) and from non-cancer female donors (*n* = 14). We isolated EVs and subjected them to our TSA assay, analyzing the expression of the nine candidate markers along with CD63 (**Fig. 7A** and **Fig. S9**). All samples showed positive CD63 values (above the control IgG level) to validate EV presence. Yet no significant difference (*P* = 0.668; unpaired two-sided *t*-test) was observed between the non-cancer and OvCa cohorts (**Fig. 7B**).

**Table 1.**
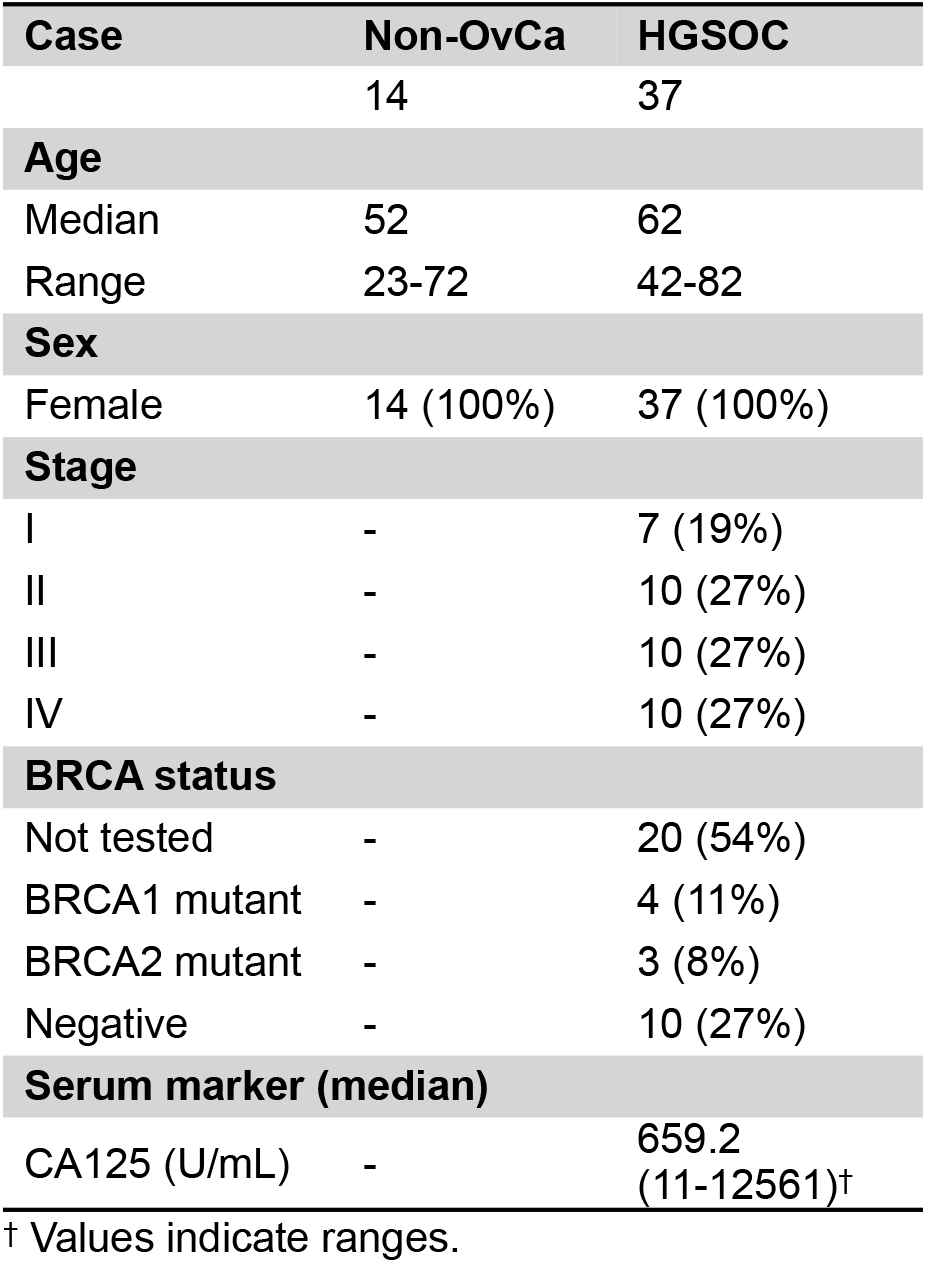
Clinical information of patients.

**Figure 7.**
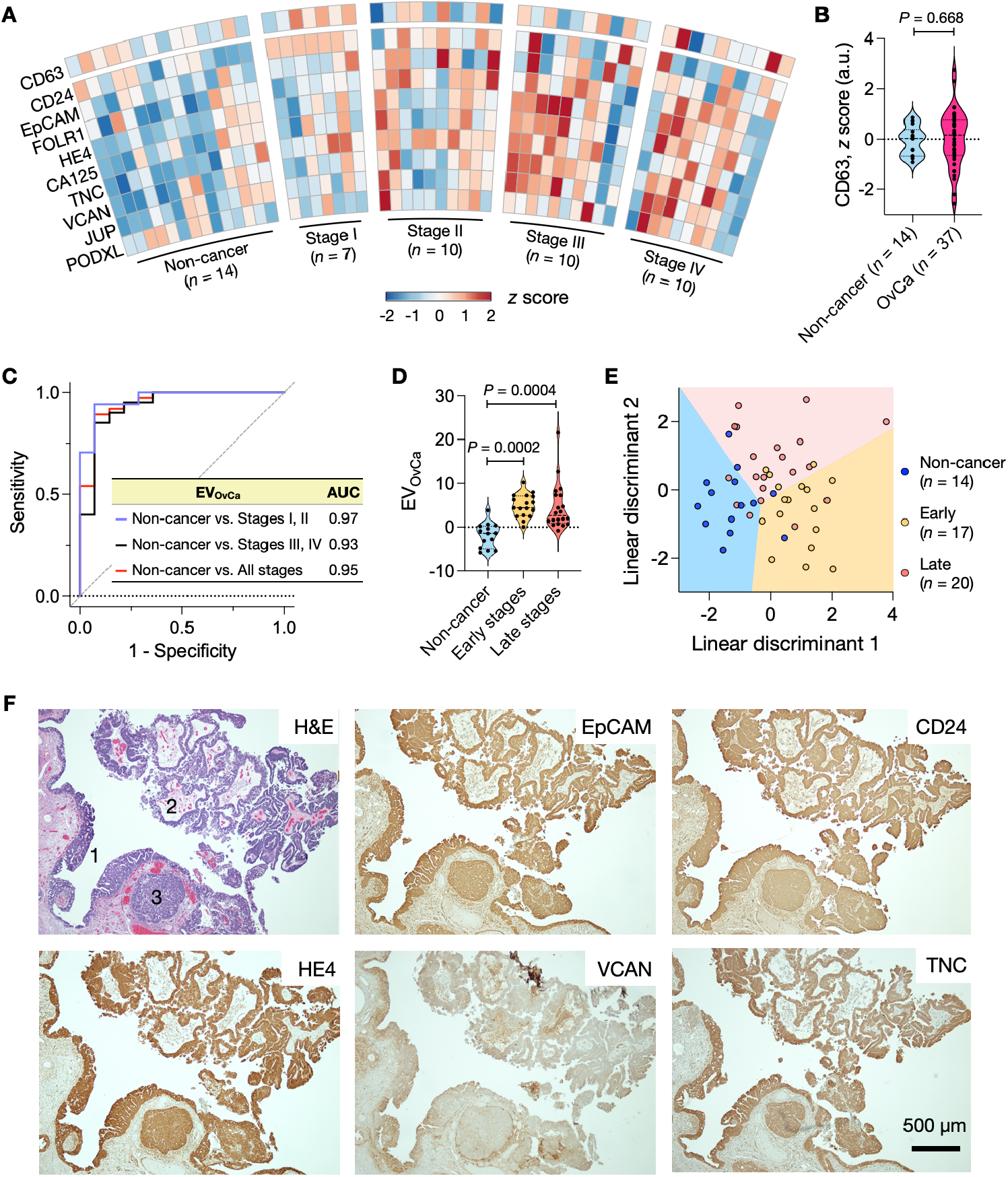
Profiling of plasma EVs from OvCa patients. **(A)** EVs from clinical plasma samples were profiled for OvCa markers and CD63 (*n* = 14, non-cancer individuals; *n* = 37, OvCa patients). The expression of each marker was normalized (z-score) and displayed in a heatmap. **(B)** CD63 measurements confirmed the presence of EVs both in non-cancer and OvCa plasma samples. No significant difference was observed in CD63 expression between the two cohorts (*P* = 0.688; non-paired, two-sided *t*-test). **(C)** Five markers (EpCAM, CD24, HE4, VCAN, TNC) were chosen from a regression analysis, and their expressions were combined to define the EV_OvCa_ score. In the receiver operating characteristic (ROC) analysis, the EV_OvCa_ score achieved high accuracies in differentiating OvCa patients from non-cancer individuals. AUC, area under the curve. **(D)** EV_OvCa_ scores were higher in early and late-stage OvCa patients than in non-cancer individuals but were similar among OvCa cohorts (Tukey’s multiple comparisons test). Early, stages I & II; Late, stages III & IV. **(E)** Linear discriminant analysis (LDA) model of top-five markers (EpCAM, CD24, HE4, VCAN, TNC) differentiated three groups: non-cancer individuals, early-stage patients, and late-stage patients. The overall classification accuracy was 74.5%. **(F)** Tumor tissue of HGSOC patients was stained positive for OvCa markers (EpCAM, CD24, HE4, VCAN, TNC). In the H&E micrograph, annotated are STIC lesions (1) and HGSOC (2 & 3). See **Fig. S12** for other OvCa markers.

We first focused on overall OvCa diagnostics through an EV protein profiling approach. We narrowed down our nine OvCa candidate markers to top-five proteins (i.e., EpCAM, CD24, HE4, VCAN, TNC) *via* lasso (least absolute shrinkage and selection operator) analyses. We then defined an EV_OvCa_ score by combining the expression of these five markers through logistic regression (see **Methods** for detail). The EV_OvCa_ score achieved high accuracy for OvCa detection (sensitivity, 0.89; specificity, 0.93) with an area under the curve (AUC) of 0.95 (**Fig. 7C**). The EV_OvCa_ score also outperformed single markers and even the nine-marker combination (see **Table S1** and **Fig. S10** for comparison). We further compared EV_OvCa_ scores among three groups: non-cancer individuals (*n* = 14), early-stage (I & II) HGSOC (*n* = 17), and late-stage (III & IV) HGSOC (*n* = 20). The average EV_OvCa_ scores from both HGSOC groups were significantly higher (*P* < 0.0002) than the non-cancer group’s average (**Fig. 7D** and **Fig. S11**) but were similar between them (*P* = 0.949; all Tukey’s multiple comparisons tests).

We next explored whether the nine OvCa markers can simultaneously classify the three groups (i.e., non-cancer, early-stage HGSOC, and late-stage HGSOC). We applied linear discriminant analysis (LDA) for this multi-class separation while varying the number of markers used as predictors. The five markers (EpCAM, CD24, HE4, VCAN, TNC) from OvCa diagnostics were selected again (**Table S2**), achieving the highest classification accuracy while minimizing the number of markers. The overall accuracy of the three-group classification was 0.75 with a 95% confidence interval of (0.63, 0.89).

Importantly, the five-marker model differentiated early-stage HGSOC group from both late-stage HGSOC and non-cancer groups (**Fig. 7E** and **Table S3**) with a specificity of 0.91 (= 31/34) and a sensitivity of 0.76 (= 13/17). We further validated our EV results by examining the tumor tissues of OvCa patients (*n* = 8). The STIC lesions and early HGSOCs were stained positive for the five markers, with strong expression in both STIC and HGSOC (**Fig. 7F** and **Fig. S12**).

## DISCUSSION

The success of nascent or future molecular tests for early OvCa detection will require high standards to be met or exceeded. Putative biomarkers should be highly specific to minimize false positives, sensing modalities should be tuned to catch weak signals from small tumors, and test procedures/assays should be affordable and minimally invasive for translation into a population-wide screening. The current study was designed to systematically address these challenges, with a focus on HGSOC, the most common and lethal OvCa subtype. We reasoned that EVs could be a potent analytic target since they are avidly produced by expanding tumor cells and readily accessible in peripheral blood. We defined HGSOC markers by analyzing EVs derived from oncogenic murine FT cells; marker expression increased in circulating EVs when HGSOC was initiated in an orthotopic murine model. To our knowledge, this is the first study to examine fallopian tube-derived EVs in the HGSOC setting. A pilot study with human patient samples further supported our preclinical findings; plasma EV analysis enabled us to identify OvCa cases (AUC of 0.97 for OvCa vs. non-cancer) and differentiate early-stage from advanced-stage cancers.

Our marker selection was guided by emerging insights that HGSOC may arise from precursor lesions within the fallopian tube. We used transformed mFT cells with a dominant negative *Tp53*^*R172H*^ mutation, loss of *Pten*, and varying *Brca* deletions. Alterations in these genes are recognized as the earliest changes seen in fallopian tube STICs and a subset of HGSOCs^6^. An average woman’s lifetime risk of developing OvCa is 1 in 78. However, this risk increases to nearly 39% in women carrying *BRCA1* mutations and up to 11% in women with *BRCA2* mutations. *TP53* mutation was found in more than 95% of HGSOC patients^49,50^. Complete and partial loss of *PTEN* was about 15% and 50-60%, respectively, in HGSOC cases^51^. The modified mFT cells harbored these common genotype variations. We selected candidate targets from mFT-EV proteomes and exploited a mouse model that recapitulated tumor initiation and progression. The EV signature of HGSOC was then identified by analyzing serial blood samples from these animals. Of note, mFT cells can also be used to model tumor metastasis and test new therapeutics, thus supporting their additional scientific and clinical potential impact^31,52–56^

Future research should interrogate EVs’ diagnostic impact for monitoring precursor evolution before or following risk reduction surgery (in high-risk contexts) or the chance encounter with an isolated STIC. First, there is a need to increase patient cohorts across the entire spectrum of the disease (i.e., benign, early, advanced stages, as well as the other more rare ovarian cancer subtypes). Such specimens will be crucial in validating the marker panel for early detection and can be used to explore which EV signatures are retained and which are specific to subtypes. We do note that our patient population reflects a single institution experience. Acquiring patient samples from multiple institutions would further enable us to account for ethnic and geographical diversities. Second, we could include other EV-associated cargo in the diagnostic algorithms. For instance, analyzing EV RNAs may boost current detection sensitivity through enzymatic target amplification and can provide complementary molecular traits (e.g., gene mutation). Herein, EV protein results could be exploited to immuno-capture early-stage OvCa EVs, thereby enhancing diagnostic specificity. Third, we can expand marker panels to obtain diverse tumor information. Previous studies have shown that EV molecular profiles can be correlated with OvCa drug resistance, poor prognostics, and metastatic propensities^57,58^. The orthotopic mFT mouse model could be an excellent translational test bed to corroborate such findings and refine markers, as it can mimic tumor initiation, progression, and spread into the peritoneal cavity. Moreover, mFT mouse models bearing various genetic subtypes have shown a potential for genotype-specific monitoring of tumor progression and treatment responses^59^. These advances should deepen our mechanistic insight into OvCa’s origin, early evolution and eventually improve patient outcomes through informed cancer management.

## MATERIALS AND METHODS

### mFT cell generation

Murine fallopian tube cells were isolated from *Brca/Tp53/Pten* GEMMs of the following four genotypes: mFT3707 (*Brca1*^*+/-*^, *Tp53*^*mut*^, *Pten*^*-/-*^), mFT3635 (*Brca1*^*-/-*^, *Tp53*^*mut*^, *Pten*^*-/-*^), mFT3665 (*Brca2*^*+/-*^, *Tp53*^*mut*^, *Pten*^*-/-*^) and mFT3666 (*Brca2*^*-/-*^, *Tp53*^*mut*^, *Pten*^*-/-*^). The fallopian tubes were extracted, digested for 48 hours in a solution of 25 mL of Minimum Essential Media supplemented with 35 mg of Pronase and 2.5 mg of DNase, and then plated in a 96-well plate. After approximately two weeks, at which point cell death became apparent, 1 µg/mL doxycycline hyclate (Sigma-Aldrich, D9891) was added to the media to trigger Cre-mediated recombination of the genes of interest under the control of the PAX8 promoter. Cells were initially propagated in 96-well plates until they were transferred to a 10 cm plate. For most cell lines, this process took roughly 10 passages and two months. The mFT culture media consisted of equal parts DMEM:F12 and M199 supplemented with HEPES pH 7.4 (10 mM), glutamine (2 mM), EGF (10 ng/mL), ITS-A (10 µg/mL), Bovine Pancreas Insulin (10 µg/mL), Hydrocortisone (0.5 µg/mL), Cholera Toxin (25 ng/mL), Retinoic Acid (25 ng/mL), BSA (1.25 mg/mL), Heat-inactivated FBS (0.75% by volume), and FBS (1% by volume).

### Luciferization of cells

mFT cells were plated and grown to 75% confluency in a 6-well plate. 0.89 μg VSV-G plasmid (Addgene, 8454), 1.38 μg Delta 8.9 plasmid, and 1.78 μg of a luc/mCherry lentiviral construct were added to 250 μL of Opti-MEM (Invitrogen, 31985070). In a separate vial, 10 μL of Lipofectamine 2000 (Invitrogen, 11668019) was added to 250 μL of Opti-MEM and incubated for 5 minutes at room temperature (RT). After incubation, the lentiviral mixture was combined with the Lipofectamine solution and incubated at RT for 20 minutes. This mixture was then added to one of the wells and incubated for 24 hours at 37 °C. The following day, the media was replaced with antibiotic-free DMEM and incubated at 37 °C for another 24 hours. On the fourth day, the media was collected and centrifuged at 1000 rpm for 10 minutes, and the supernatant was stored before refreshing the media in the plate again with antibiotic-free DMEM. The media in the other 5 wells of the plate was aspirated and replaced with 600 μL of the collected supernatant from the transfected well in addition to 8 μg/mL of polybrene and 1.4 mL of antibiotic-free DMEM. The same process was repeated the following day. After this procedure, the cells were sorted for mCherry expression at the Brigham & Women’s Flow Cytometry Core.

### Characterization of mFT cells

To analyze cell genes, we isolated genomic DNA using a Gentra Puregene Cell Kit (Qiagen, 158767) according to the manufacturer’s protocol. DNA was used to analyze recombination *via* polymerase chain reaction (PCR) as previously described ^31^. For the *Brca1* PCR, 100 ng of DNA was used for the positive control, while 500 ng was used for the heterozygous and homozygous deletion samples. For the *Brca2* PCR, 100 ng of DNA was used for the control and the heterozygote, while 1,000 ng of DNA was used for the homozygous deletion sample. For *Tp53* and *Pten* PCR, we used 300 ng of DNA samples. All gel electrophoresis was completed using a 1517bp-100bp ladder (BioLabs, N3231L). To analyze protein cellular protein, we performed western blots for CA125 (Abbiotec, 250556), γ-H2AX (CST, 9718S), PAX8 (Proteintech, 10336), and β-actin (Sigma, A2228) as previously described^31^. Western blots were completed with a 250kD-10kD Protein Precision Plus Kaleidoscope Standard (BioRad, 1610375). Gel images were taken with the ChemiDoc XRS+ System (BioRad, 1708265).

### Immunofluorescent cell imaging

We placed glass coverslips in a 6-well culture plate and soaked them with 70% ethanol for 1 hour. After washing the coverslips with PBS, we seeded about 3 × 10^5^ cells per well and cultured them overnight in culture media at 37 °C. The cells were then washed with PBS and fixed with 4% paraformaldehyde in PBS for 20 minutes at RT. The slips were then rinsed with PBS, followed by permeabilization with 0.5% Triton X-100 in PBS (PBS-T; 30 min, RT). The cells were rinsed again with PBS, and the coverslips were moved to a paraffin-covered plate that functioned as an incubation chamber. The cells were blocked with 1% BSA in 0.05% PBS-T for 1 hour at RT, using approximately 250 µL of this blocking solution per coverslip. The blocking solution was aspirated, and the cells were incubated with 200-300 µL of CA125 (Abbiotec, 250556), p53 (Leica Biosystems, CM5), Ki67 (Novus Bio, NB110-89719), or WT1 (Abcam, ab15249) antibody solution diluted in the aforementioned blocking buffer. After incubation (1 hour, RT), cells were triple-washed with 0.05% PBS-T in 5-min intervals. Finally, cells were incubated with fluorophore-conjugated secondary antibodies (1 hour, RT) and then triple-washed with 0.05% PBS-T in 5-min intervals. The coverslips were mounted onto slides using Fluoromount-G, allowed to dry, sealed, and imaged using an EVOS FL Auto 2 microscope. We used ImageJ (version 10.2) to adjust for background fluorescence.

### Spheroid formation

We seeded about 6.5 × 10^3^ cells per well in a 96-well ultra-low adhesion plate (Corning, CLS3474). The cells were imaged on days 3, 7, and 14. For spheroid-to-monolayer experiments, 6 × 10^5^ cells were seeded per well in a 6-well ultra-low adhesion plate (Corning, CLS3471) prior to plating in a 10 cm dish.

### EV proteomics

We plated an equal number of mFT cells (mFT3635, mFT3666) in a 150 cm^2^ culture dish. When cells reached about 99% confluency, we rinsed them with PBS and allowed them to grow in in serum-depleted culture medium (48 hours). We next collected cell culture media and removed cell debris *via* centrifugation (10,000 ×g, 3 min). To collect EVs, we first concentrated supernatants using a centrifugal filter (Centricon Plus-70, Millipore Sigma) and loaded a concentrated media (0.5 mL) to a qEV column (IZON, SP1). EV fractions (F7-F9) were collected (1.5 mL). We measured total protein concentrations using a Qubit protein assay kit (ThermoFisher, Q33212). For proteome profiling, isolated EVs were sent to an external vendor (BGI company) that performed EV lysis, protein recovery, sample digestion, and nano-flow LC-MS/MS analysis. Abundant proteins were selected by the spectral counting method provided by BGI.

### Nanoparticle tracking analysis

We used NanoSight LM10 (Malvern) equipped with a 405 nm laser. Samples were diluted in PBS to obtain the recommended particle concentration (25-100 particles/ frame). For each sample, we recorded three 30-second video clips and analyzed them using NTA software (version 3.2). All the measurements were performed under the same acquisition settings (e.g., camera level, detection threshold).

### Single EV imaging

We diluted plasma EVs in PBS and removed aggregated using a 0.22-µm Millex-GV syringe filter (Millipore Sigma, SLGV004SL). Filtered EVs were captured on a glass slide (Electron Microscopy Sciences). Following a 30-min incubation at RT, the slide was twice washed with PBS. After incubation with a mixture of a fixation buffer (4% formaldehyde) and a permeabilization buffer (BD Biosciences, 554723), EV samples were treated with Superblock (ThermoFisher, 37518) for 1 hour and labeled (90 min, RT) with a mixture of primary antibodies against CD63, CD9, PAX8, and one of the mFT markers (CA125, VCAN, TNC, FOLR1, HE4). IgG antibody was used as a control. Samples were then triple-washed with 0.01% PBS-Tw (PBS buffer containing 0.01% Tween 20), labeled (30 min, RT) with fluorophore-conjugated secondary antibodies, and triple-washed again with PBS-Tw. We finally took fluorescence images using a Nikon A1R confocal microscope with a 60 × objective. All the acquisition settings (i.e., objective, image size, laser power, gain) were kept the same. Images were processed (i.e., background subtraction) and analyzed with Comdet v0.5.5 in Image J software (NIH).

### High-throughput EV analysis

We suspended EV isolates in 0.05 M carbonate-bicarbonate buffer (pH, 9.4; Millipore Sigma, C3041) and loaded the solution (25 µL per well) into a high-binding 384-well plate (Greiner, 781077). The plate was placed on an orbital shaker, and EVs were allowed to adsorb to the plate surface (overnight, 4 °C). Following the incubation, we washed the plate with PBS buffer containing 0.1% Tween 20 (0.1% PBS-Tw) two times and treated it with a blocking buffer (2% BSA in PBS) for 1 hour at RT. After washing the plate with 0.1% PBS-Tw, we added primary antibodies (1% BSA in 0.1% PBS-Tw), allowed them to react (1 hour, RT), and removed excess antibodies *via* triple-washing with 0.1% PBS-Tw. Subsequently, EVs were similarly labeled with biotinylated secondary antibodies (1% BSA in 0.1% PBS-Tw). We next reacted samples with streptavidin-HRP (ThermoFisher, 21130; 1% BSA in 0.1% PBS-Tw) for 30 min at RT, triple-washed them with 0.1% PBS-Tw, and performed tyramide signal amplification by adding 6 µg/mL tyramide-biotin (Millipore Sigma, SML2135) in amplification buffer (0.003% Hydrogen peroxide in 0.1× borate buffer). After a 30-min incubation at RT, we triple-washed samples with 0.1% PBS-Tw and labeled them with streptavidin-BV510 (Biolegend, 405234; 1% BSA in 0.1% PBS-Tw). After a 30-min incubation, samples were washed (0.1% PBS-Tw2), and their fluorescence intensities were measured by a plate reader (Tecan) at 405/490 excitation/emission wavelengths with 20 nm bandwidth.

### Tumor mouse model

Animal studies were conducted following the guidelines provided by the Institutional Animal Care and Use Committee (IACUC) of Brigham and Women’s Hospital, Harvard Medical School, and the Whitehead Institute at the Massachusetts Institute of Technology. For orthotopic injections, we anesthetized 12-16 weeks old NOD.Cg-*Prkdc*^*scid*^ *Il2rg*^*tm1Wjl*^*/*SzJ (NOD-*scid* IL2Rgamma^null^) female mice (Whitehead Institute) with 1.25% Avertin *via* intraperitoneal injection (400 mg/kg). Using aseptic techniques, we made a 1 cm incision along the thoracic region between the third and fourth mammary gland (roughly 0.5 cm from the spine) to expose the peritoneum and another small incision (0.5 cm in length) into the peritoneal wall above the ovary and ovarian fat depot. The adipose tissue and ovary were pulled through the peritoneal incision to expose 0.5 cm of adipose tissue. We injected about 10^6^ mFT cells (resuspended in 10-20 µL of a 1:1 dilution of Matrigel in PBS) into the fallopian tube using a Hamilton syringe. Once the procedure was complete, we carefully pushed back the adipose depot through the peritoneal wall and closed the peritoneum using sterile biodegradable J204G 4-0 Vicryl violet sutures (Ethicon). After suturing the peritoneum, we gathered the skin and closed the incision with wound clips. Mice were kept warm on a heating pad until fully recovered. Meloxicam was administered intraperitoneally (5 mg/kg) immediately after surgery and 24 hours later. For intraperitoneal injections, 6-week-old NOD.CB17-Prkdcscid/NCrCrl (SCID) mice (Charles River Laboratories, Strain #394) were injected intraperitoneally with 5 × 10^6^ cells resuspended in PBS. For bioluminescence imaging, mice were injected (IP) with luciferin (25% of their body weight) in PBS. After 4 minutes, isoflurane was administered to anesthetize the mice for imaging on the Xenogen IVIS-100 Imaging System.

### Serial EV monitoring from tumor-bearing mice

About 200 µL of blood was sampled from the retroorbital sinus of a mouse using a heparinized microhematocrit tube. Blood samples were immediately centrifuged at 2000 × g for 15 minutes at 4°C, plasma supernatant was collected, and plasma samples were stored at -80°C. To isolate EVs, we processed about 50 µL of plasmas with qEV columns (IZON, SP2) per the manufacturer’s instruction.

### Human samples

The study protocol was reviewed and approved by the Institutional Review Board of Dana–Farber/Harvard Cancer Center (IRB number, 07-049). Informed consent was obtained from all participants of this clinical study. Plasma samples were centrifuged at 2000 ×g for 3 min to remove cell debris, and supernatants were collected. About 100 µL of plasmas were used for EV isolation through qEV columns (IZON, SP2). We analyzed EVs for protein markers following the high-throughput EV analysis protocol.

### Immunohistochemistry

We collected tumors and other organs (ovary, fallopian tube, uterus) from mice for immunohistochemistry (IHC). IHC samples were sent to the Specialized Histopathology Core at Harvard Medical School for staining. Additionally, some IHC experiments were performed at the Whitehead Institute, Massachusetts Institute of Technology. For ascites IHC samples, ascites were collected from the mice, spun down, and cultured under the same conditions as their mFT cell line origins. After the ascites lines were established, we collected about 2 × 10^7^ cells *via* centrifugation, fixed the cells in 10% formalin, and embedded them in paraffin. The cells were then sectioned and adhered to slides and stained by the Specialized Histopathology Core at Harvard Medical School. Slides were stained for PAX8 (ProteinTech, 10336), p53 (Abcam, 1431), STMN1 (CST, 13655), and WT1 (Sigma, 348M-9).

### Biomarker IHC validation

The study protocol was reviewed and approved by the Institutional Review Board of Massachusetts General Hospital (IRB protocol, 2022P001059). Formallin-fixed paraffin-embedded sections of human patient samples were baked at 56 °C for 30 min and deparaffinized with 10-min xylene washes (two times). Samples were then rehydrated with 5-min washes of ethanol at increasing dilutions (100%, 100%, 95%, 75%). Endogenous peroxidase was then blocked using a 15-min incubation in a 1:1 solution of 100% ethanol and 3% hydrogen peroxide. Antigen retrieval was performed by boiling samples in citrate buffer (Dako, S1699) for 15 min at 110 °C using a NxGen Decloaking Chamber (Biocare, DC2012). Slides were then washed with PBS and blocked with 10% goat serum for an hour at RT. After blocking with serum, the samples were blocked for endogenous biotin and avidin binding sites following manufacturer instructions (Vector, SP-2001). Sections were then incubated overnight at 4 °C with their respective primary antibodies diluted in PBS. The following day, samples were washed in tris-buffered saline supplemented with 0.05% tween-20 (TBS-Tw) prior to incubating with biotin-conjugated secondary antibodies for one hour at RT. Slides were then washed again with TBS-Tw and incubated for an additional hour at RT with HRP-conjugated streptavidin (Vector, SA-5004; 1:100). After a final set of TBS-Tw washes, slides were visualized using DAB, counterstained with hematoxylin, air-dried overnight, and coverslips were mounted using Permount mounting medium (Fisher SP15-500).

### Statistical analysis

We performed statistical analysis using GraphPad Prism version 9.4 (GraphPad Software Inc.) or R version 4.1.0. For all statistical tests, *P* values < 0.05 were considered significant. (i) Marker selection for OvCa detection. We selected candidate markers from the EV profiling data by applying the lasso regression. Specifically, we calculated the cross-validation error (CVE) and determined the tuning parameter (λ) that minimized CVE. We used the glmnet package in R. (ii) OvCa diagnosis. We assessed the diagnostic ability of EV markers by conducting a ROC analysis. Individual EV markers and their combinations were considered. For each biomarker combination, optimal weights were determined *via* logistic regression. We constructed ROC curves and determined level cutoffs that maximized the sum of sensitivity and specificity. Standard formulas were used to define sensitivity (true positive rate), specificity (true negative rate), and accuracy [(true positive + true negative)/(positive + negative)]. AUCs were compared following Delong’s method. EV_OvCa_ had the highest AUC value with the minimal number of EV markers used. Analyses were performed using the pROC package in R. (iii) Linear discriminant analysis (LDA). We constructed an LDA model to classify clinical samples into three groups: non-cancer, early-stage OvCa (stages I & II), and late-stage OvCa (stages III & IV). We varied the number of EV markers used as predictors and constructed a confusion matrix. The five-marker (EpCAM, CD24, HE4, VCAN, TNC) LDA model showed the highest classification accuracy with the following linear discriminants: LD1 = 0.165·EpCAM + 0.638·CD24 – 0.191·HE4 + 0.172·VCAN + 1.126·TNC; LD2 = 0.268·EpCAM + 0.175·CD24 + 1.106·HE4 + 0.495·VCAN – 0.922·TNC. Confidence intervals for the accuracy of each model were estimated using the bootstrap method with 1000 replicates. We used the MASS and the Boot packages in R.

### Antibodies

All antibodies with their associated dilution and application can be found in **Tables S4, S5, and S6**.

## Data availability

The main data supporting the results of this study are available within the paper and its supplementary information. The raw patient datasets generated and analyzed during the study are available from the corresponding authors on reasonable request, subject to approval from the Institutional Review Board of Massachusetts General Hospital.

## Special thanks

Christopher R. Getchell (BWH), Vivian L. Zhang (BWH), Abigail Cohen (BWH), Morgan Krush (BWH), and Dominique Zarrella (MGH).

