## supplementary information for "Profiling extracellular vesicles in circulation enables the early detection of ovarian cancer"

[#]Equal contribution (J.E.M., S.I., A.W.O., E.T.M.)

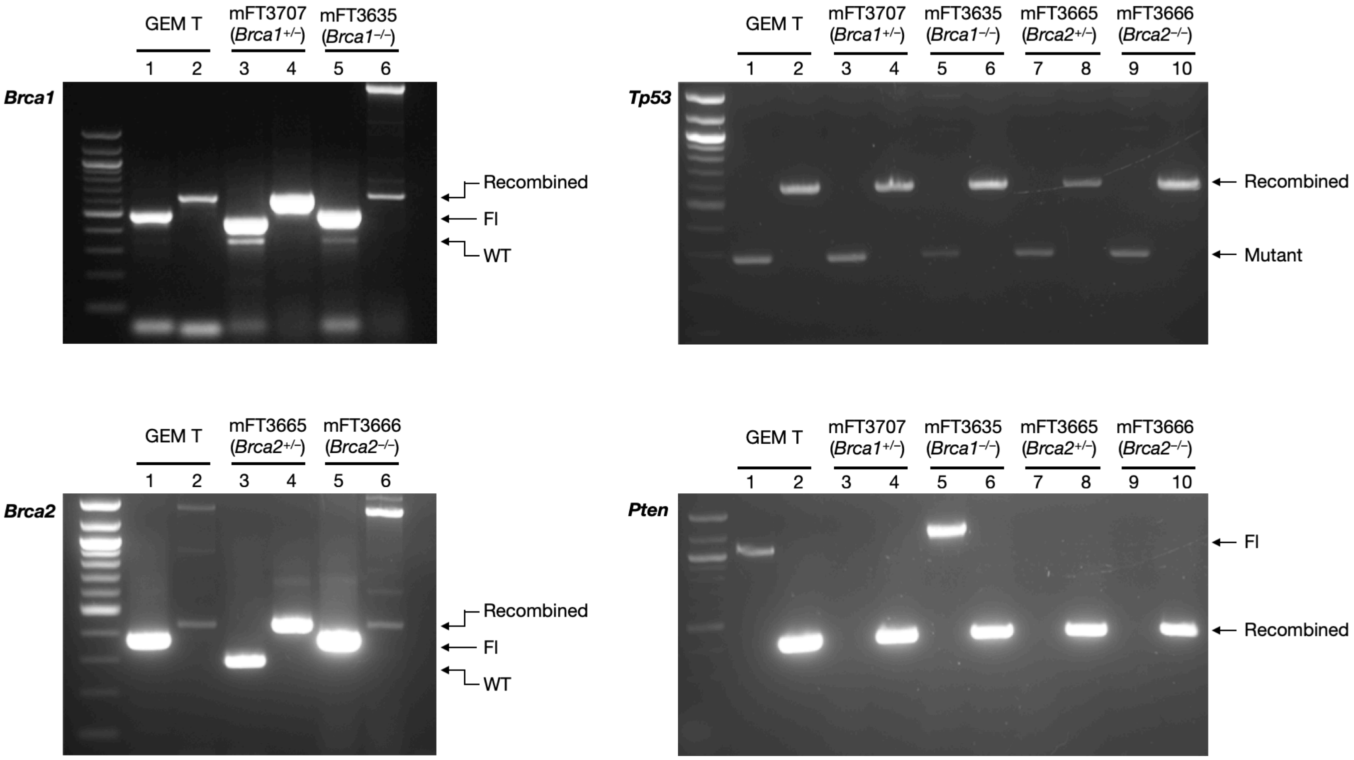


**Supplementary Figure S1. Genotyping of mFT cell lines.** *Brca1*, *Brca2*, *Tp53*, and *Pten* genes were amplified by a polymerase chain reaction and measured by gel electrophoresis. The following cell lines were characterized: mFT3707 ($Brca1^{+/-}$, $Tp53^{mut}$, $Pten^{-/-}$), mFT3635 ($Brca1^{-/-}$, $Tp53^{mut}$, $Pten^{-/-}$), mFT3665 ($Brca2^{+/-}$, $Tp53^{mut}$, $Pten^{-/-}$) and mFT3666 ($Brca2^{-/-}$, $Tp53^{mut}$, $Pten^{-/-}$). Odd numbered lanes have unrecombined DNA, and even numbered lanes have *Cre*-mediated recombined DNA. GEM T, tumors from genetically engineered model tumors; Fl, floxed allele; WT, wild-type allele.

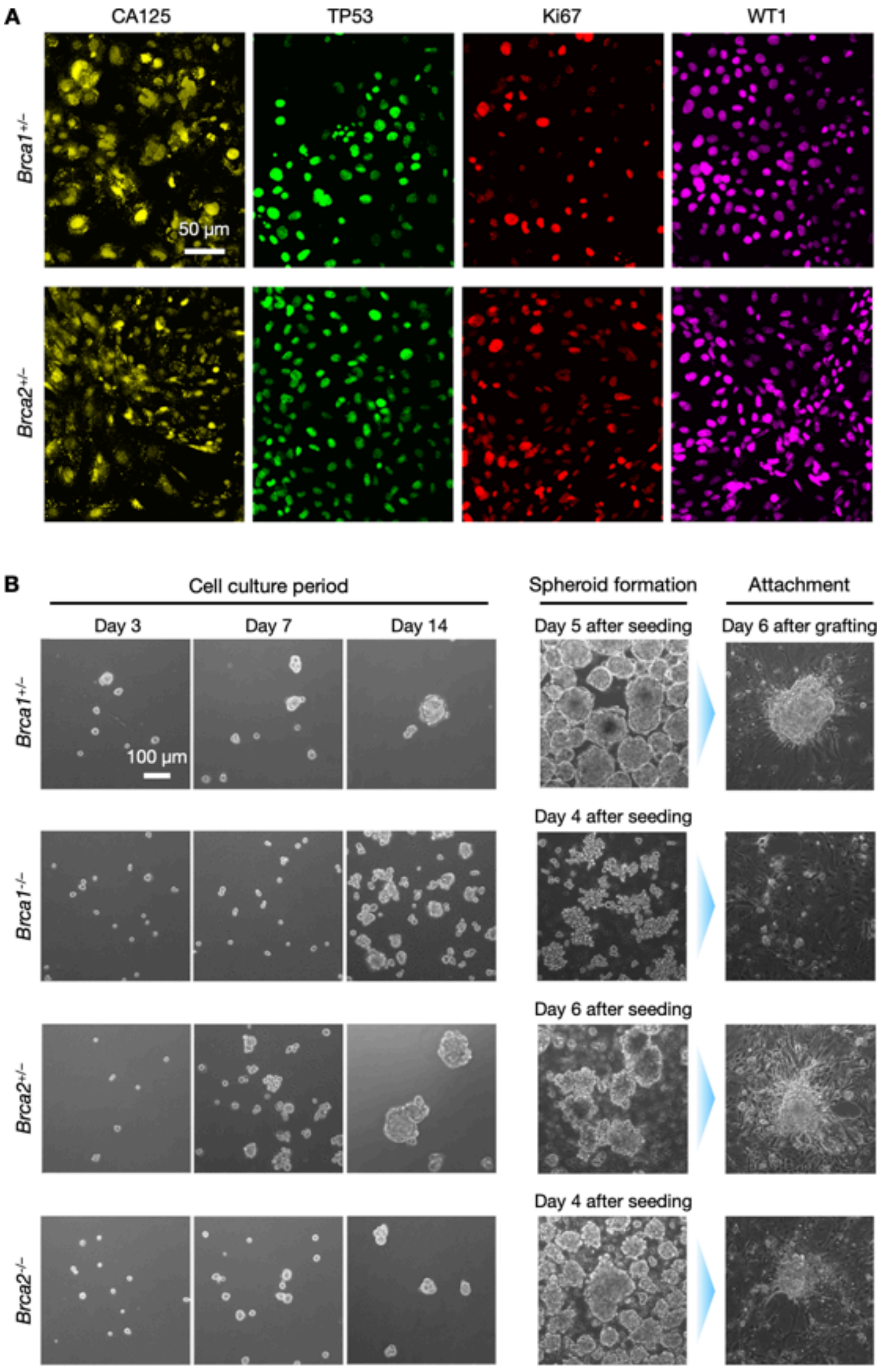


**Supplementary Figure S2. *In vitro* Characterization of mFT cell lines. (A)** Immunofluorescence staining of CA125, TP53, Ki67, and WT1 in additional mFT cell lines: mFT3707 (*Brca1*$^{+/-}$) and mFT3665 (*Brca2*$^{+/-}$). **(B)** Spheroid formation of mFT cell lines. Microscope images show spheroid formation and growth over a 14-day period. Spheroids were then transferred from ultra-low adhesion plates to attachment plates to demonstrate spheroids' capability to adhere to surfaces.

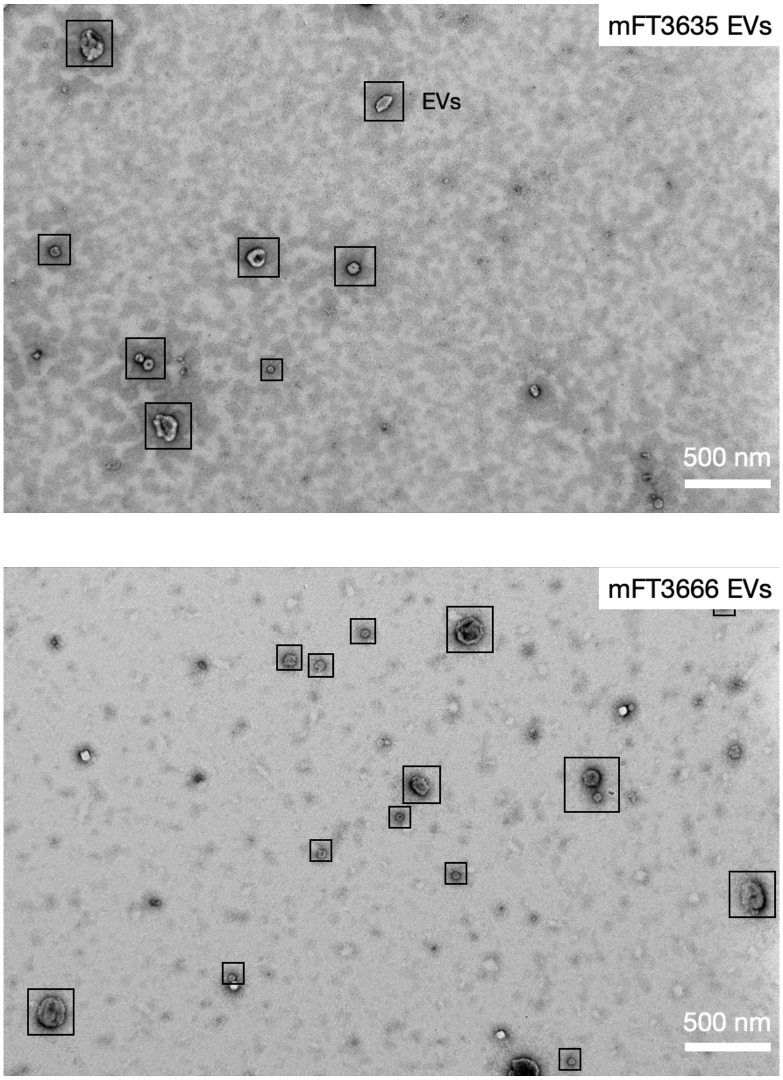


**Supplementary Figure S3. Large field-of-view TEM images of mFT EVs.** (Top) EVs isolated from mFT3635 cells. (Bottom) EVs isolated from mFT3666 cells. Black squares indicate EVs.

**A**

| Cell line | EV sample replicate | Input [EV] (particles/mL) | Number of proteins identified |
|---|---|---|---|
| mFT3635 (*Brca1*$^{-/-}$) | 1 | $2.5 \times 10^{11}$ | 571 |
| | 2 | $9.1 \times 10^{10}$ | 507 |
| | 3 | $1.7 \times 10^{10}$ | 465 |
| mFT3666 (*Brca2*$^{-/-}$) | 1 | $1.4 \times 10^{9}$ | 133 |
| | 2 | $6.3 \times 10^{9}$ | 177 |
| | 3 | $4.0 \times 10^{8}$ | 157 |

**B**

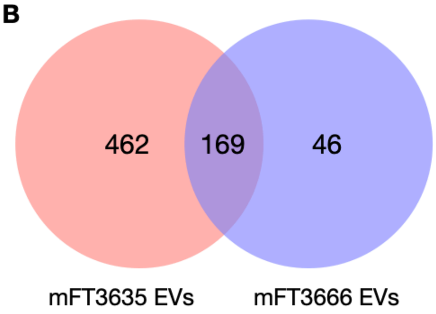


**Supplementary Figure S4. Proteome analysis of mFT EVs. (A)** Summary characteristics of mFT-EV samples isolated from mFT3635 and mFT3666 cell lines. Three replicates from each cell line were used for the proteome profiling. **(B)** The Venn diagram shows the distribution of proteins identified in mFT3635- and mFT3666-EV samples.

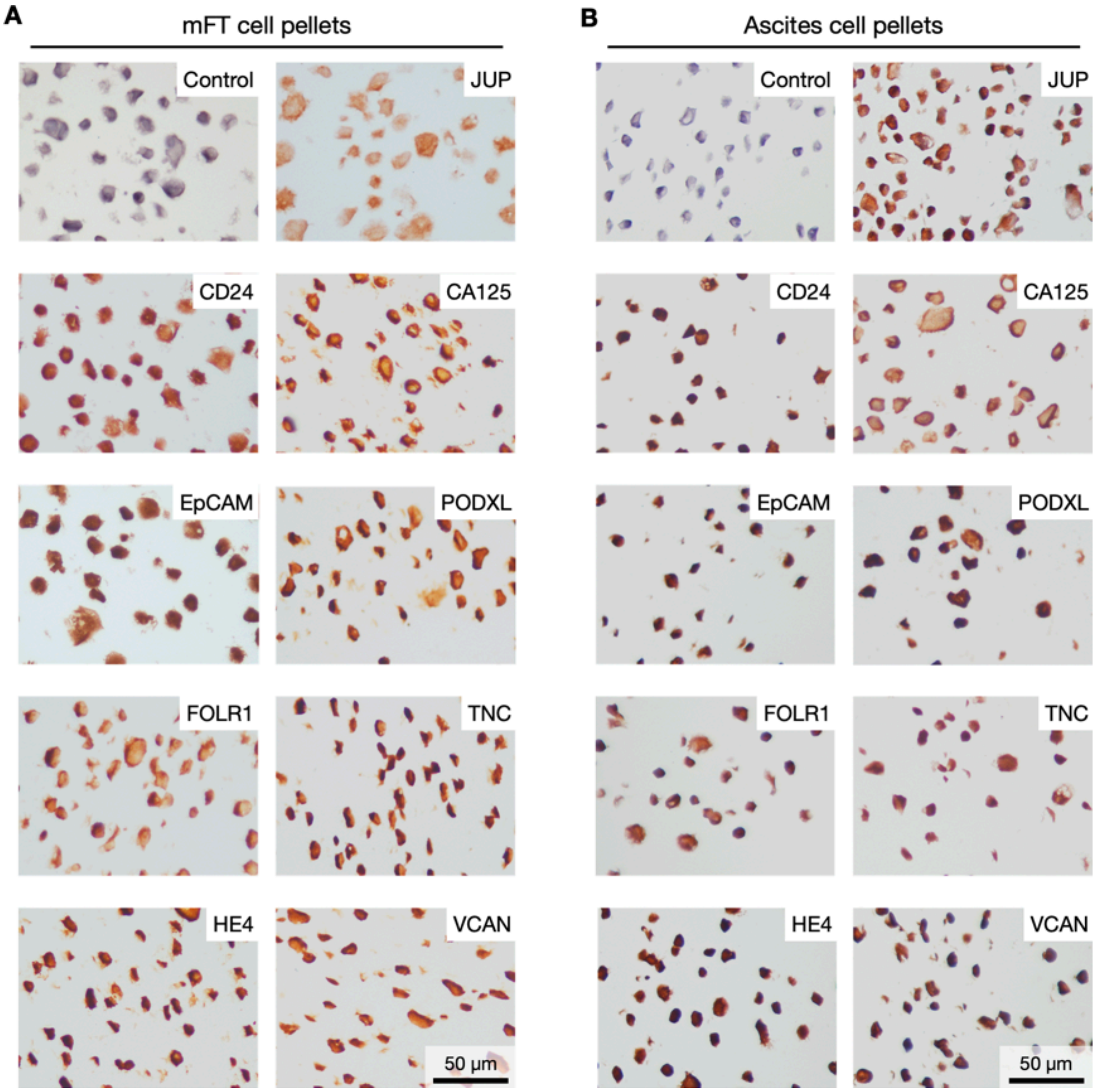


**Supplementary Figure S5. Expression of candidate markers in mFT parent and ascites cells. (A)** IHC staining of mFT3666 parental cell lines. (**B**) IHC staining of ascites cells collected from a mouse implanted with mFT3666 cells.

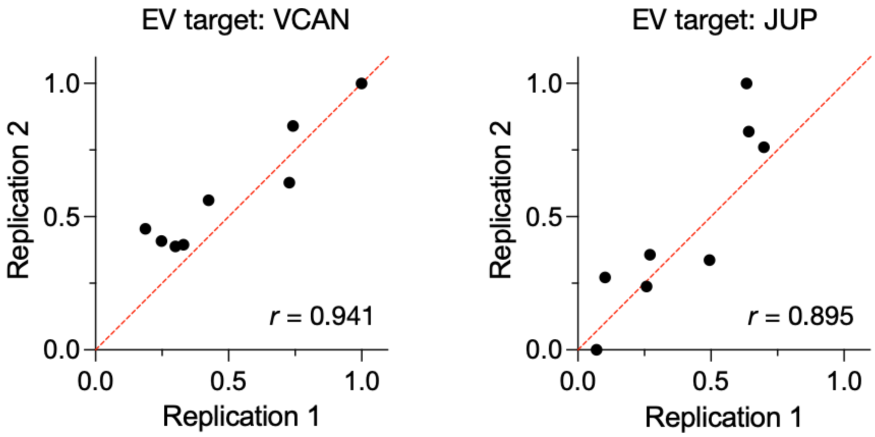


**Supplementary Figure S6. Reproducibility of the tyramine application assay.** Eight plasma EV samples were assayed for VCAN (left) and JUP (right) expression on two different days (replications 1 and 2). For a given marker, a high correlation was observed between two independent measurements. Pearson's *r* values were 0.941 (VCAN; *P* = 0.005) and 0.895 (JUP; *P* = 0.0027). The dotted line is a reference line (slope = 1). Pearson's r values were determined using GraphPad Prism version 9.4.

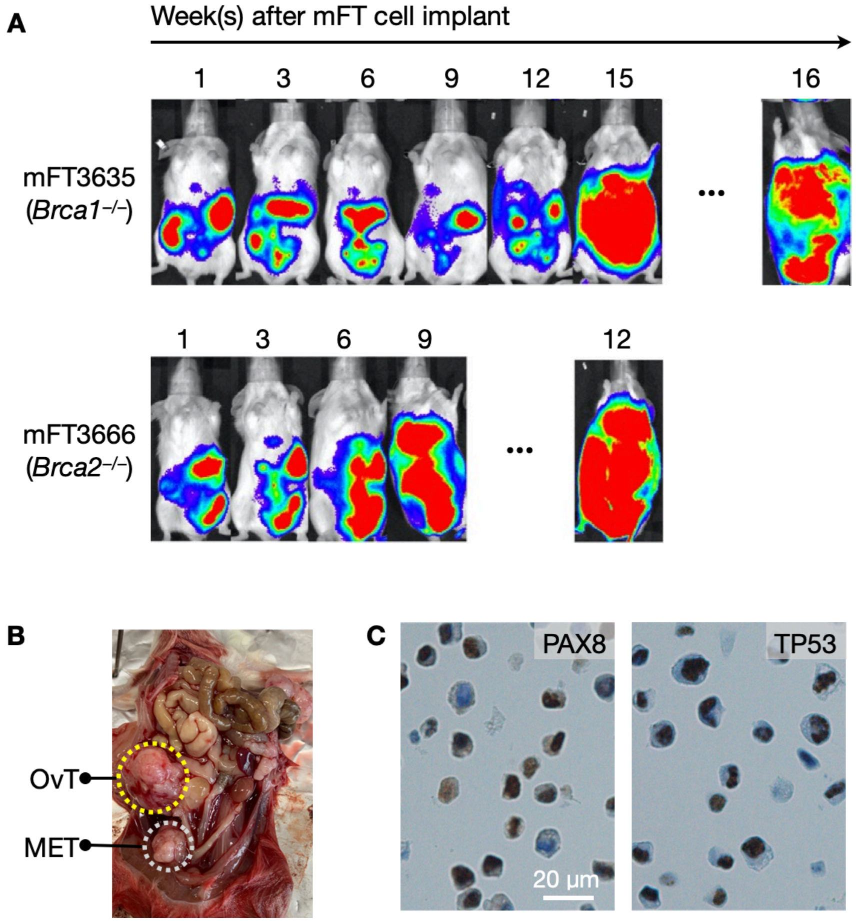


**Supplementary Figure S7. Tumorigenic properties of oncogenic mFT cell lines. (A)** NOD SCID mice were implanted IP with oncogenic mFT cells expressing luciferase. Bioluminescence imaging confirmed the spread of the tumor throughout the peritoneum. **(B)** Presence of tumors in an NSG mouse orthotopically implanted with mFT3666 cells. OvT, primary tumor; MET, metastasis. **(C)** Representative images of ascites cells from mFT3666 engrafted mice. Cells were stained positive for PAX8 and TP53.

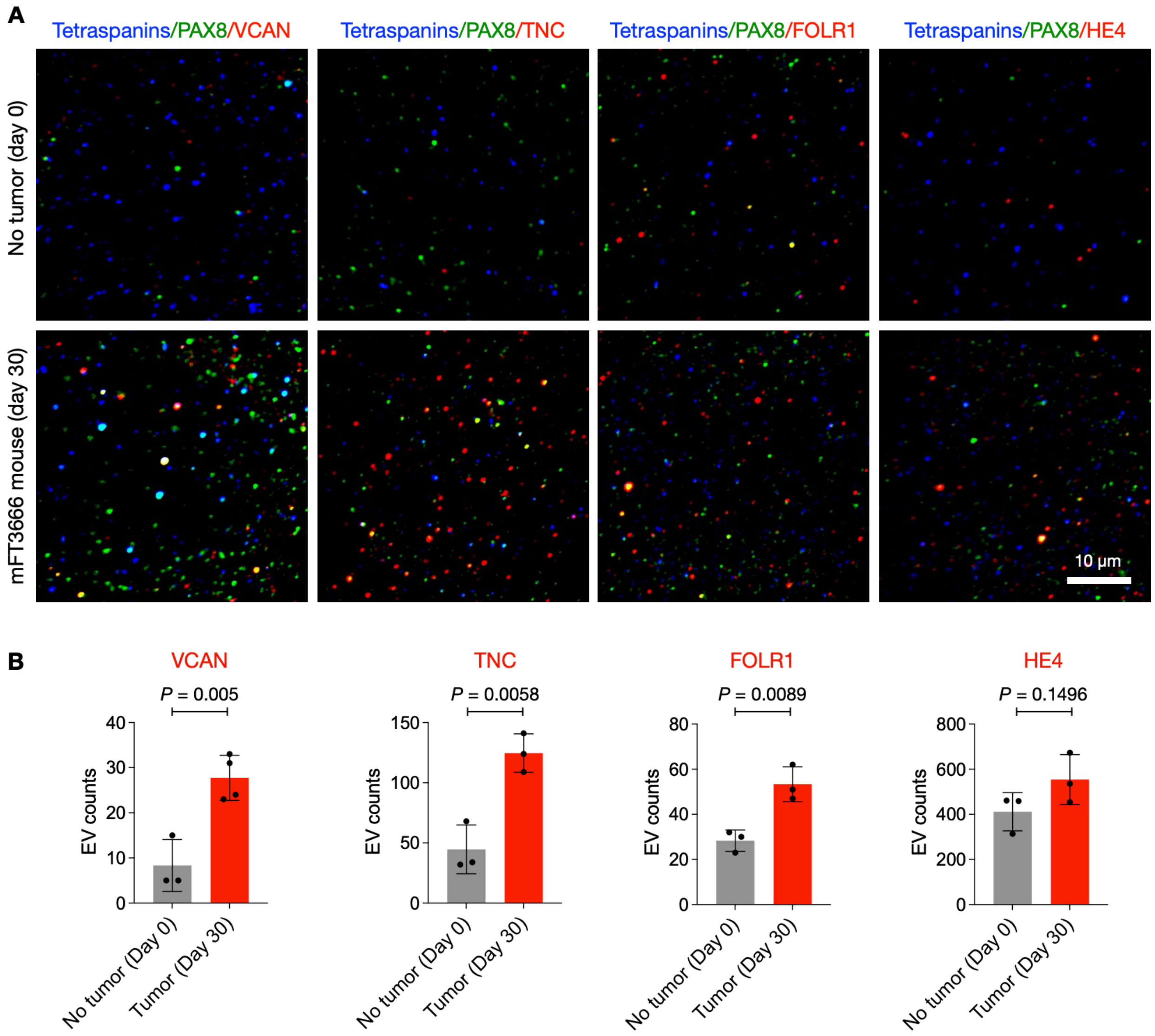


**Supplementary Figure S8. Single EV analysis of murine plasma EVs. (A)** Plasma samples were obtained from an animal before the mFT cell (*Brca2*$^{-/-}$) implant (day 0) and during tumor growth (day 30). EVs were immunostained for tetraspanins (CD63, CD9, CD81), PAX8 (FT epithelial marker), and OvCa markers. **(B)** More plasma EVs were counted positive for OvCa markers in the day-30 sample. Each dot in the graph represents the marker-positive EV number in a given field of view.

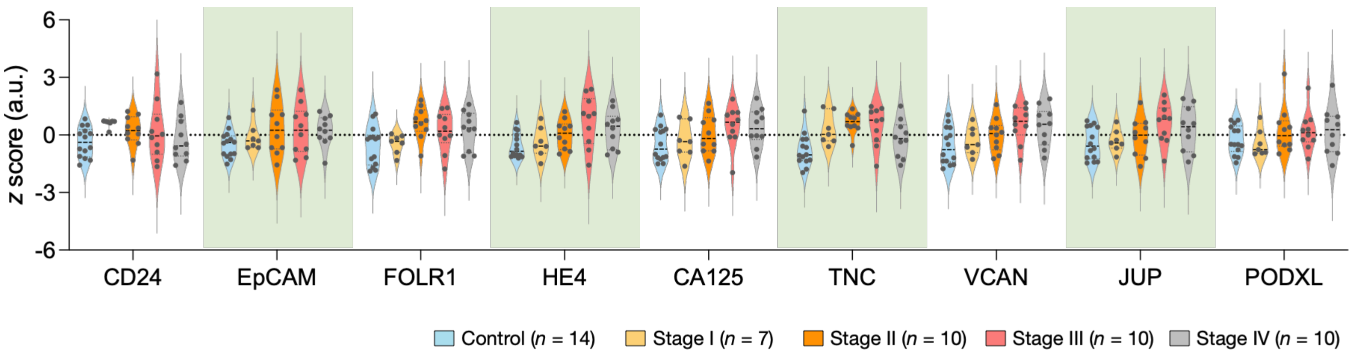


**Supplementary Figure S9. Expression levels of nine OvCa markers.** EVs in clinical samples were profiled. For each marker, z-scores were calculated and stratified by the tumor stage.

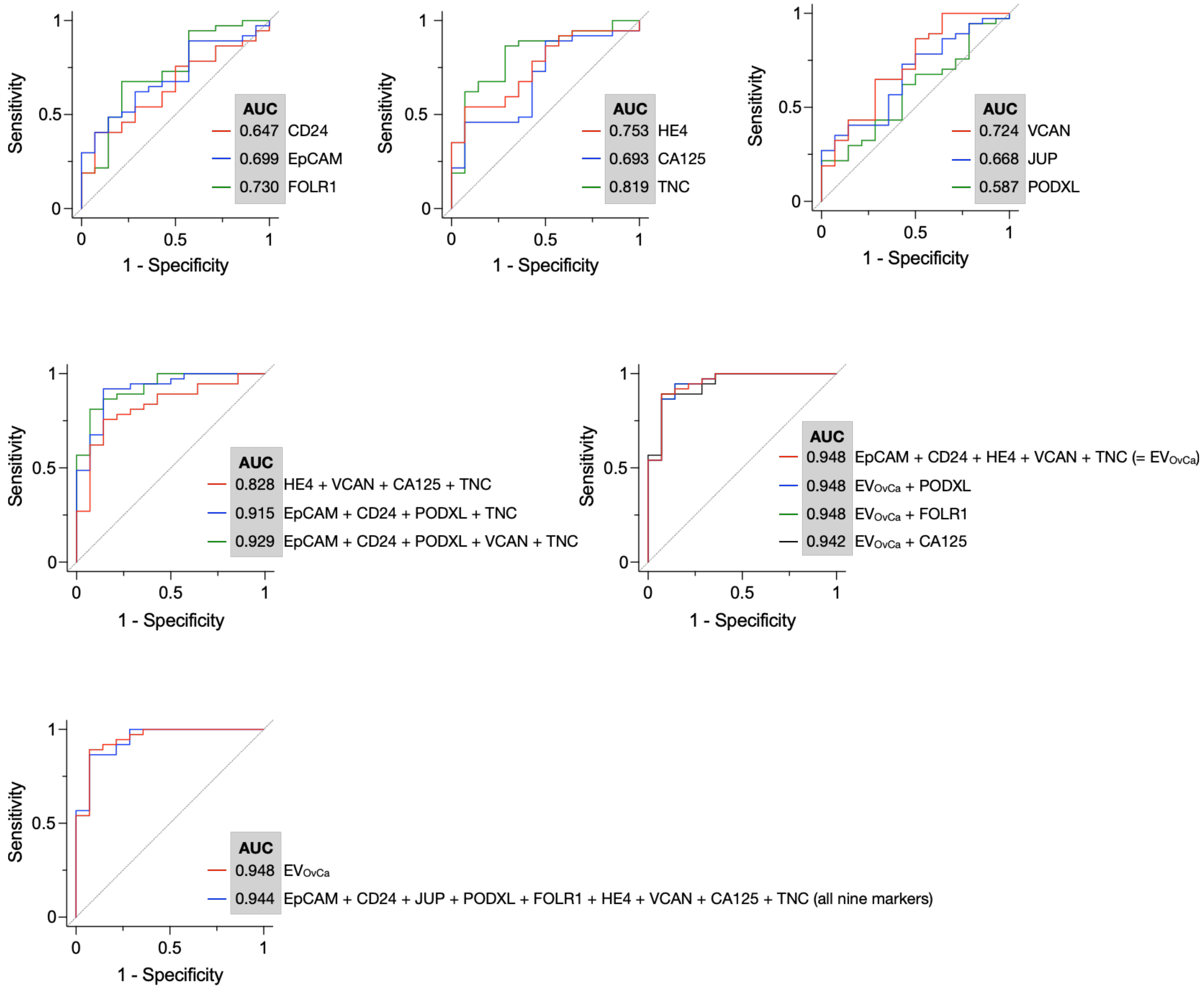


**Supplementary Figure S10. ROC analyses for OvCa detection**. Single markers and their combinations were used as classifiers for control (*n* = 14) vs. cancer (*n* = 37) cases. EV$_{OvCa}$ achieved the highest area under the curve (AUC) with a minimal marker set.

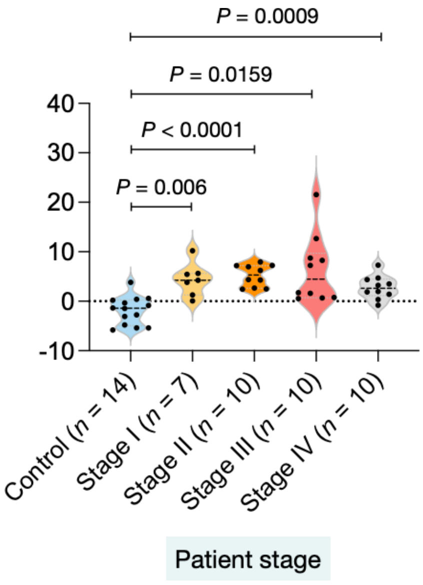


**Supplementary Figure S11. Stratified $EV_{OvCa}$ scores per tumor stage.** Adjusted *P* values were obtained from Dunnett's T3 multiple comparisons test.

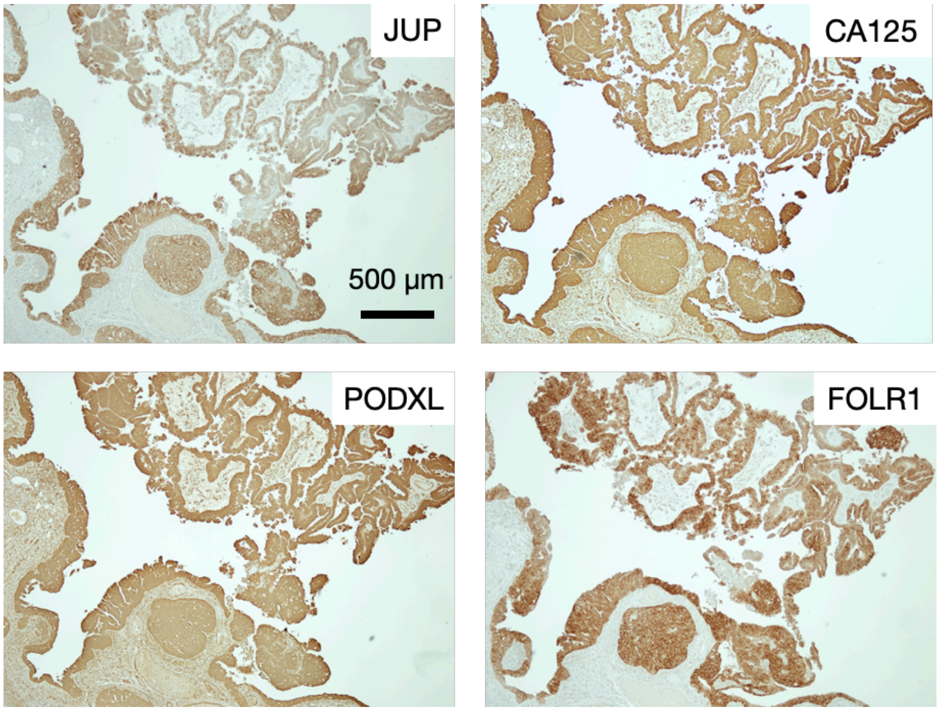


**Supplementary Figure S12. Images of IHC stained tissue from an OvCa patient.** STIC and HGSOC were stained positive for JUP, CA125, PODXL, and FOLR1.

**Supplementary Table 1. Statistics of OvCa diagnoses.**

| Markers | Control vs. early stage | | | | Control vs. all stage | | | |
|---|---|---|---|---|---|---|---|---|
| | AUC | Sen | Spe | Acc | AUC | Sen | Spe | Acc |
| CD24 | 0.782 | 0.965 | 0.714 | 0.742 | 0.647 | 0.405 | 0.857 | 0.549 |
| EpCAM | 0.672 | 0.941 | 0.429 | 0.710 | 0.699 | 0.487 | 0.857 | 0.588 |
| FOLR1 | 0.714 | 0.647 | 0.786 | 0.710 | 0.730 | 0.676 | 0.786 | 0.706 |
| HE4 | 0.693 | 0.941 | 0.429 | 0.710 | 0.753 | 0.541 | 0.929 | 0.647 |
| CA125 | 0.626 | 0.882 | 0.500 | 0.710 | 0.693 | 0.892 | 0.500 | 0.765 |
| TNC | 0.882 | 1.000 | 0.643 | 0.839 | 0.819 | 0.865 | 0.714 | 0.824 |
| VCAN | 0.639 | 1.000 | 0.357 | 0.710 | 0.724 | 0.865 | 0.500 | 0.765 |
| JUP | 0.576 | 0.765 | 0.500 | 0.645 | 0.668 | 0.730 | 0.571 | 0.686 |
| PODXL | 0.542 | 1.000 | 0.214 | 0.645 | 0.587 | 0.216 | 100.000 | 0.431 |
| HE4 + VCAN + CA125 + TNC | 0.891 | 0.765 | 0.929 | 0.839 | 0.828 | 0.757 | 0.857 | 0.765 |
| EpCAM + CD24 + PODXL + TNC | 0.971 | 1.000 | 0.857 | 0.935 | 0.915 | 0.919 | 0.857 | 0.882 |
| EpCAM + CD24 + PODXL + VCAN + TNC | 0.979 | 0.941 | 0.929 | 0.935 | 0.929 | 0.811 | 0.929 | 0.824 |
| EpCAM + CD24 + HE4 + VCAN + TNC (= $EV_{OvCa}$)† | 0.966 | 0.941 | 0.929 | 0.935 | 0.948 | 0.892 | 0.929 | 0.902 |
| EpCAM + CD24 + HE4 + VCAN + TNC + PODXL | 0.966 | 0.941 | 0.929 | 0.935 | 0.948 | 0.946 | 0.857 | 0.882 |
| EpCAM + CD24 + HE4 + VCAN + TNC + FOLR1 | 0.966 | 0.941 | 0.929 | 0.935 | 0.948 | 0.946 | 0.857 | 0.902 |
| EpCAM + CD24 + HE4 + VCAN + TNC + CA125 | 0.966 | 0.941 | 0.929 | 0.935 | 0.942 | 0.892 | 0.929 | 0.863 |
| EpCAM + CD24 + JUP + PODXL + FOLR1 + HE4 + VCAN + CA125 + TNC | 0.966 | 0.941 | 0.929 | 0.935 | 0.944 | 0.865 | 0.929 | 0.902 |

† The minimal marker set (in yellow) was selected for the highest diagnostic accuracy. AUC, area under the curve; Sen, sensitivity; Spe, specificity; Acc, accuracy.

**Supplementary Table 2. Performance summary of linear discriminant analysis (LDA).**

| Markers | Correctly predicted cases | | | | |
|---|---|---|---|---|---|
| | Non-cancer (*n* = 14) | Early (*n* = 17) | Late (*n* = 20) | Overall accuracy | 95% confidence interval |
| CD24 | 2 | 9 | 10 | 0.412 | (0.255, 0.628) |
| EpCAM | 6 | 0 | 15 | 0.412 | (0.314, 0.608) |
| FOLR1 | 6 | 0 | 15 | 0.412 | (0.294, 0.588) |
| HE4 | 8 | 6 | 13 | 0.529 | (0.353, 0.647) |
| CA125 | 7 | 4 | 17 | 0.549 | (0.353, 0.686) |
| TNC | 9 | 7 | 8 | 0.471 | (0.353, 0.647) |
| VCAN | 7 | 4 | 15 | 0.510 | (0.353, 0.647) |
| JUP | 5 | 7 | 13 | 0.490 | (0.333, 0.628) |
| PODXL | 3 | 6 | 15 | 0.471 | (0.294, 0.607) |
| HE4, VCAN, CA125, TNC | 8 | 11 | 13 | 0.627 | (0.529, 0.824) |
| EpCAM, CD24, PODXL, TNC | 12 | 9 | 10 | 0.608 | (0.529, 0.843) |
| EpCAM, CD24, PODXL, VCAN, TNC | 10 | 13 | 13 | 0.706 | (0.588, 0.863) |
| EpCAM, CD24, HE4, VCAN, TNC | 10 | 13 | 15 | 0.745† | (0.628, 0.882) |
| EpCAM, CD24, HE4, VCAN, TNC, PODXL | 10 | 13 | 15 | 0.745 | (0.648, 0.902) |
| EpCAM, CD24, HE4, VCAN, TNC, FOLR1 | 10 | 12 | 15 | 0.725 | (0.628, 0.902) |
| EpCAM, CD24, HE4, VCAN, TNC CA125 | 10 | 12 | 14 | 0.706 | (0.647, 0.902) |
| EpCAM, CD24, JUP, PODXL, FOLR1, HE4, VCAN, CA125, TNC | 10 | 14 | 14 | 0.745 | (0.706, 0.922) |

† The minimal marker set for LDA (in yellow) was selected for the highest accuracy in the three-group classification.

**Supplementary Table 3. Confusion matrix of the 5-marker LDA model.**

| | | Clinical diagnosis | | |
|---|---|---|---|---|
| | | Control | Early OvCa | Late OvCa |
| LDA prediciton | Control | 10 | 0 | 3 |
| | Early OvCa | 1 | 13 | 2 |
| | Late OvCa | 3 | 4 | 15 |

**Supplementary Table 4. Primary antibodies used in the study.**

| Target | Application | Dilution | Vendor | Catalog # |
|---|---|---|---|---|
| CA125 | WB | 1:1000 | Abbiotec | 250556 |
| γ-H2AX | WB | 1:1000 | CST | 9718S |
| PAX8 | WB | 1:1000 | ProteinTech | 10336-1-AP |
| β-actin | WB | 1:2000 | Sigma | A2228 |
| CA125 | IF | 1:250 | Abbiotec | 250556 |
| p53 | IF | 1:100 | Leica | CM5 |
| KI-67 | IF | 1:100 | Novus | NB110-89719 |
| WT-1 | IF | 1:300 | Abcam | ab15249 |
| PAX8 | Murine IHC | 1:600 | ProteinTech | 10336-1-AP |
| p53 | Murine IHC | 1:200 | Abcam | ab1431 |
| Stathim | Murine IHC | 1:2000 | CST | 13655 |
| WT-1 | Murine IHC | 1:100 | Sigma | 348M-9 |
| CD24 | Murine IHC | 1:50 | Bioss | 4891R |
| EpCAM | Murine IHC | 1:50 | Bioss | 1513R |
| FOLR1 | Murine IHC | 1:50 | Abcam | ab67422 |
| HE4 | Murine IHC | 1:50 | LsBio | LS-C409035 |
| JUP | Murine IHC | 1:50 | Genetex | GTX114156 |
| CA125 | Murine IHC | 1:50 | LsBio | LS-C408274 |
| PODXL | Murine IHC | 1:50 | Bioss | 1345R |
| TNC | Murine IHC | 1:50 | LsBio | LS-C413317 |
| VCAN | Murine IHC | 1:50 | Bioss | 2533R |
| CD24 | Human IHC | 1:250 | Novus | NBP1-46390 |
| EpCAM | Human IHC | 1:100 | Bioss | 1513R |
| FOLR1 | Human IHC | 1:100 | Abcam | ab67422 |
| HE4 | Human IHC | 1:100 | LsBio | LS-C409035 |
| JUP | Human IHC | 1:100 | Genetex | GTX114156 |
| CA125 | Human IHC | 1:100 | LsBio | LS-C408274 |
| PODXL | Human IHC | 1:100 | Bioss | 1345R |
| TNC | Human IHC | 1:100 | LsBio | LS-C413317 |
| VCAN | Human IHC | 1:100 | Novus | NBP1-85432 |

WB, western blotting; IF, immunofluorescence; IHC, immunohistochemistry.

**Supplementary Table 5. Antibodies used for EV ELISA.**

| Target | Species | Dilution | Vendor | Catalog # | Isotype |
|---|---|---|---|---|---|
| CD24 | Mouse | 1:100 | Biolegend | 101801 | Rat IgG2b |
| CD24 | Human | 1:250 | Biolegend | 311102 | Mouse IgG2a |
| EpCAM | Mouse/Human | 1:250, 1:40 | Invitrogen | MA5-12436 | Mouse IgG1 |
| FOLR1 | Mouse/Human | 1:500, 1:200 | LsBio | LS-C119975 | Rabbit IgG polyclonal |
| HE4 | Mouse/Human | 1:750, 1:300 | LsBio | LS-C409035 | Rabbit IgG polyclonal |
| JUP | Mouse/Human | 1:250, 1:100 | LsBio | LS-B2204 | Goat IgG polyclonal |
| CA125 | Mouse/Human | 1:1000, 1:400 | LsBio | LS-C408274 | Rabbit IgG polyclonal |
| PODXL | Mouse | 1:200 | R&D systems | MAB1556 | Rat IgG2b |
| PODXL | Human | 1:500 | R&D systems | AF1658 | Goat IgG polyclonal |
| TNC | Mouse/Human | 1:500, 1:200 | LsBio | LS-C413317 | Rabbit IgG polyclonal |
| VCAN | Mouse | 1:400 | Bioss | BS-2533R | Rabbit IgG polyclonal |
| VCAN | Human | 1:200 | Millipore Sigma | HPA004726 | Rabbit IgG polyclonal |
| CD9 | Mouse | 1:100 | Biolegend | 124802 | Rat IgG2a |
| CD9 | Human | 1:250 | BD Biosciences | 555370 | Mouse IgG1 |
| CD63 | Mouse | 1:200 | Biolegend | 143902 | Rat IgG2a |
| CD63 | Human | 1:500 | Ancell | 215-820 | Mouse IgG1 |
| CD81 | Mouse | 1:200 | Biolegend | 104901 | Armenian Hamster IgG1 |
| TSG101 | Mouse | 1:200 | Genetex | 118736 | Rabbit IgG polyclonal |
| Histone 2B | Mouse | 1:200 | Biolegend | 606302 | Rat IgG2a |
| CD63-biotin | Mouse | 1:200 | Biolegend | 143918 | - |

| IgG isotype | Species | Dilution | Vendor | Catalog # |
|---|---|---|---|---|
| Rat IgG2a | Mouse/Human | *a* | Biolegend | 400502 |
| Rat IgG2b | Mouse/Human | *a* | Biolegend | 400602 |
| Mouse IgG1 | Mouse/Human | *a* | Biolegend | 400102 |
| Mouse IgG2a | Mouse/Human | *a* | Biolegend | 400202 |
| Rabbit polyclonal IgG | Mouse/Human | *a* | Biolegend | 910801 |
| Goat IgG Isotype | Mouse/Human | *a* | Novus Biologicals | NB410-28088 |
| Armenian Hamster IgG | Mouse/Human | *a* | Biolegend | 400902 |

*a*. IgG isotype controls were diluted to the same concentration of primary antibody for each biomarker

| Biotinylated secondary | Species | Dilution | Vendor | Catalog # |
|---|---|---|---|---|
| Goat Anti-Rat IgG, Biotin-SP | Mouse/Human | 1:750 | Millipore Sigma | AP183B |
| Goat Anti-Rabbit IgG, Biotin-SP | Mouse/Human | 1:500 | Millipore Sigma | AP132B |
| Goat Anti-Mouse IgG, Biotin-SP | Mouse/Human | 1:500 | Millipore Sigma | AP124B |
| Goat Anti-Armenian hamster IgG, Biotin | Mouse/Human | 1:250 | Biolegend | 405501 |
| Rabbit Anti-Goat IgG, Biotin | Mouse/Human | 1:750 | ThermoFisher | A16146 |

**Supplementary Table 6. Antibodies used for single EV imaging.**

| Primary antibody | Dilution | Vendor | Catalog # | Isotype |
|---|---|---|---|---|
| CD9 | 1:50 | Biolegend | 124802 | Rat IgG2a |
| CD63 | 1:50 | Biolegend | 143902 | Rat IgG2a |
| PAX8 | 1:50 | ProteinTech | 60145-4-Ig | Mouse IgG |
| CA125 | 1:100 | LsBio | LS-C408274 | Rabbit IgG polyclonal |
| VCAN | 1:100 | Bioss | BS-2533R | Rabbit IgG polyclonal |
| TNC | 1:100 | LsBio | LS-C413317 | Rabbit IgG polyclonal |
| FOLR1 | 1:100 | LsBio | LS-C119975 | Rabbit IgG polyclonal |
| HE4 | 1:100 | LsBio | LS-C409035 | Rabbit IgG polyclonal |
| **Secondary antibody** | | | | |
| Anti-Rat AlexaFluor647 | 1:400 | Abcam | ab150155 | - |
| Anti-Mouse AlexaFluor488 | 1:400 | Abcam | ab150105 | - |
| Anti-Rabbit AlexaFluor594 | 1:400 | Abcam | ab150076 | - |
